# Unveiling Tissue Structure and Tumor Microenvironment from Spatial Omics by Hypergraph Learning

**DOI:** 10.1101/2024.05.15.594168

**Authors:** Yi Liao, Chong Zhang, Zhikang Wang, Fei Qi, Weitian Huang, Shangyan Cai, Junyu Li, Jiazhou Chen, Robin B. Gasser, Zhiyuan Yuan, Jiangning Song, Hongmin Cai

**Affiliations:** School of Computer Science and Engineering, South China University of Technology, Guangzhou, Guangdong, 510006, China; Biomedicine Discovery Institute and Department of Biochemistry and Molecular Biology, Monash University, Melbourne, VIC, 3800, Australia; Monash AI Institute, Monash University, Melbourne, VIC, 3800, Australia; Faculty of Science and Technology, Beijing Normal University-Hong Kong Baptist University United International College, Zhuhai, Guangdong, 519087, China; Department of Veterinary Biosciences, Melbourne Veterinary School, Faculty of Science, The University of Melbourne, Parkville, VIC 3010, Australia; Institute of Science and Technology for Brain-Inspired Intelligence, MOE Key Laboratory of Computational Neuroscience and Brain-Inspired Intelligence, MOE Frontiers Center for Brain Science, Fudan University, Shanghai, 200433, China; Center for Medical Research and Innovation, Shanghai Pudong Hospital, Fudan University Pudong Medical Center, Fudan University, Shanghai, 201399, China

**Keywords:** Spatial omics, Spatial domain, Resolution, Hypergraph learning, Higher-order relationship

## Abstract

Spatial omics technologies have revolutionized life sciences by enabling the simultaneous acquisition of biomolecular and spatial information. Identifying spatial patterns is crucial for understanding organ development and tumor microenvironments. However, the emergence of diverse spatial omics resolutions in these technologies has made it challenging to accurately characterize spatial domains at finer resolutions. To address this, we propose HyperSTAR, a hypergraph-based method designed to precisely identify spatial domains across varying resolutions by leveraging higher-order relationships among spatially adjacent tissue programs. Specifically, a gene expression-guided hyperedge decomposition module is introduced to refine the hypergraph structure to accurately delineate spatial domains boundaries. Additionally, a hypergraph attention convolutional neural network is designed to adaptively learn the importance of each hyperedge, enhancing the model’s ability to capture complex higher-order relationships within spatially neighboring multi-spots and/or single cells. HyperSTAR outperforms existing graph neural network models in tasks such as uncovering tissue substructures, inferring spatiotemporal patterns, and denoising spatially resolved gene expressions. It effectively handles diverse spatial omics data types and scales seamlessly to large datasets. The method successfully reveals spatial heterogeneity in breast cancer sections, with findings validated through functional and survival analyses of independent clinical data. HyperSTAR represents a significant advancement in spatial omics analysis, representing a robust tool for exploring complex spatial patterns across varying resolutions and data types. Its ability to capture intricate higher-order relationships among spatially neighboring spots/cells makes it an invaluable tool for advancing research in life sciences, particularly in cancer and developmental biology.

## Introduction

In living organisms, biological process happens within a spatial context. The location of each cell is as crucial as its inherent characteristics in determining cell types, cell states and functions [1]. Single-cell omics quantifies genomes, epigenomes, transcriptomes and proteomes at the single-cell level [2–3]. Nevertheless, the process of single-cell dissociation during sample preparation results in the loss of spatial information [4]. Emerging spatially resolved transcriptomics (SRT) have facilitated the profiling of gene expression while preserving spatial location in tissues, offering valuable opportunities in exploring intricate mechanisms within biological systems [5].

Widely used SRT can be broadly categorized into two groups: (1) imaging-based approaches, and (2) next-generation sequencing (NGS)-based approaches [6–10]. These diverse technologies simultaneously capture both the gene expression and spatial location, while differing from gene throughput, sensitivity, resolution, and field of view (fov). Imaging-based methods provide subcellular resolution with high sensitivity but limited gene throughout [11–17], while NGS-based technologies generate diverse spatial resolutions, ranging from multi-cellular to subcellular levels for the captured spots with whole-transcriptome coverage but lower sensitivity [18–20].

Several computational methods have been developed to identify spatial domains in SRT data. Earlier works utilized traditional methods (e.g. K-means [21–22], Louvain [23]), which did not take into account the spatial information of SRT data, leading to a discontinuous division of regions. Recent methods have introduced spatial information to enhance spatial domain identification. For example, BayesSpace [24] employed a Bayesian statistical approach, using the information from spatial neighborhoods for spatial clustering. Deep learning methods have also been developed to integrate spatial information and gene expression of SRT data. StLearn [25] defined morphological distances based on features extracted from histology images and utilized these distances, along with spatial neighbors to smooth gene expressions. SpaGCN [26] also used histology images by employing graph convolutional networks to integrate gene expression, spatial location and image. SpaceFlow [27] utilized spatially regularized deep graph networks to integrate both expression similarity and spatial information. STAGATE [28] applied a graph attention auto-encoder to learn low-dimensional latent embeddings of SRT profiles. Although these methods have shown significant improvements on some datasets, they primarily focus on the cell-cell pairwise relationships, neglecting the higher-order structures inherent in multicellular programs. Furthermore, they fail to account for the delineation of domain boundaries, a crucial aspect for enhancing spatial domain results, as they predominantly focused on clustering. This limitation becomes more pronounced when confronted with higher spatial resolutions, making it challenging to elucidate the tissue structure comprehensively. Moreover, these approaches encounter difficulties in extending their capabilities to accommodate other omics.

To tackle these issues, we propose HyperSTAR, the first hypergraph learning-based spatial clustering method by explicitly learning the higher order tissue structures. HyperSTAR can effectively leverage complex relationships among spatially adjacent multi-cellular programs. A gene expression-guided hyperedge decomposition module and a hypergraph attention convolutional layer are employed to precisely delineate boundaries of spatial domains. Extensive experiments show HyperSTAR could reveal tissue structure and tumor spatial heterogeneity across varying spatial resolutions, demonstrating effectiveness at diverse downstream analyses, such as spatiotemporal trajectories inference, gene expression denoising and spatially variable genes (SVGs) detection.

## Results

### Overview of HyperSTAR

Spatial resolved transcriptomics (SRT) data typically contains a gene expression matrix and a spatial coordinate matrix, as illustrated in Fig. 1a. To comprehensively capture the intricate associations among spatially adjacent multiple spots, we develop a hypergraph-based method, as illustrated in Fig. 1b and Fig. 1c. The construction of this hypergraph can be primarily divided into two key steps: (1) acquiring spot representations and (2) establishing the hypergraph structure. Spot representations are acquired from pre-processed gene expression data. The hypergraph structure is attained through two stages: (1) constructing a naive hyperedge set among spatially neighboring multi-spots, and (2) conducting a gene expression-guided hyperedge decomposition module to refine these naive hyperedges. Finally, a hypergraph structure that fuses spatial information and gene expression is obtained (Fig. 1b). The initially built naive hyperedge set contains complex high-order relationships among spatially neighboring multi-spots. However, it is important to note that adjacent spots in space do not always belong to the same domain. For instance, although spots near a domain boundary are spatially adjacent, they might actually belong to different domains. To tackle this issue, a gene expression-guided hyperedge decomposition module is conducted to enhance the effects of gene expression. After constructing the hypergraph, we utilize hypergraph attention convolutional neural networks to learn low-dimensional latent embeddings of the SRT data. Subsequently, the latent embeddings are used to visualize data, identify spatial domains, infer trajectories, detect spatially variable genes (SVGs), and denoise spatial gene expression (Fig. 1c). Using the above strategy to integrate spatial information and gene expression in SRT data, HyperSTAR improves the performance of spatial omics analysis.

**Fig. 1.**
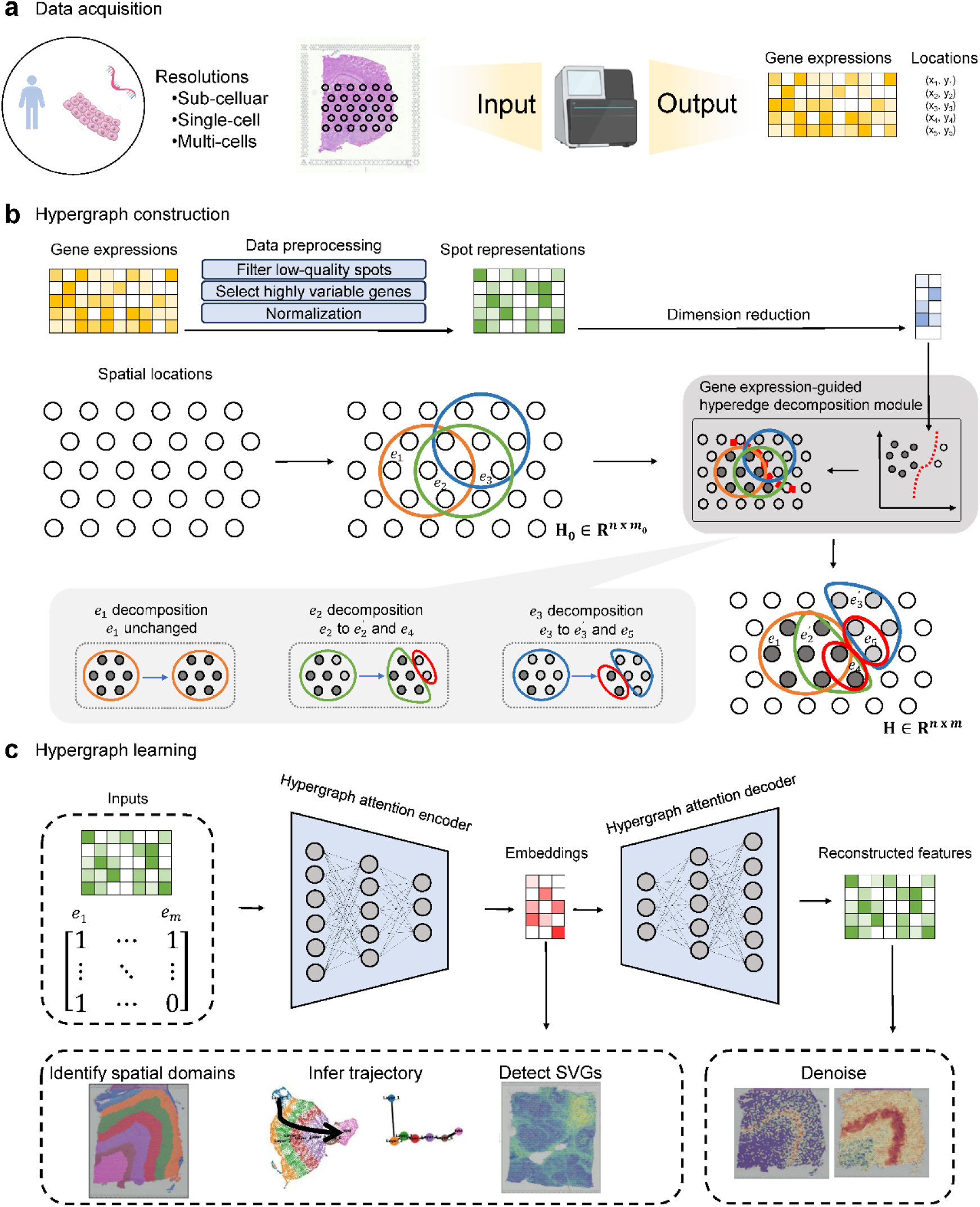
Overview of HyperSTAR. **a** Spatial resolved transcriptomics (SRT) data varies in resolution depending on the sequencing technology. Typically, SRT data consists of gene expression profiles and their corresponding spatial coordinates. **b** A hypergraph is constructed to model relationships among multi-spots. Spot representations are derived directly from gene expression data. The hypergraph structure is built in two stages: (1) for each spot, a hyperedge is formed by connecting its spatially adjacent multi-spots, and (2) hyperedge decomposition is performed to refine the hyperedges based on gene expression clustering results. **c** The refined hyperedge set and spot representations are input into hypergraph attention convolutional neural networks to learn latent embeddings of spots for downstream analyses.

### HyperSTAR achieves state-of-the-art performance on the benchmark dataset

To evaluate the performance of HyperSTAR, we first used the widely recognized SpatialLIBD dataset, a 10x Visium dataset of 12 human dorsolateral prefrontal cortex (DLPFC) sections [29]. This dataset includes manually annotated DLPFC layers (Layer 1 to Layer 6) and white matter (WM), which are defined based on morphological features and marker genes. Using these annotations as the ground truth, we employed multiple established metrics—Adjusted Rand Index (ARI), Normalized Mutual Information (NMI), Moran’I and Geary’s C—to quantitatively assess the accuracy and robustness of spatial domain identification (Fig. 2a and Supplementary Fig. S19).

**Fig. 2.**
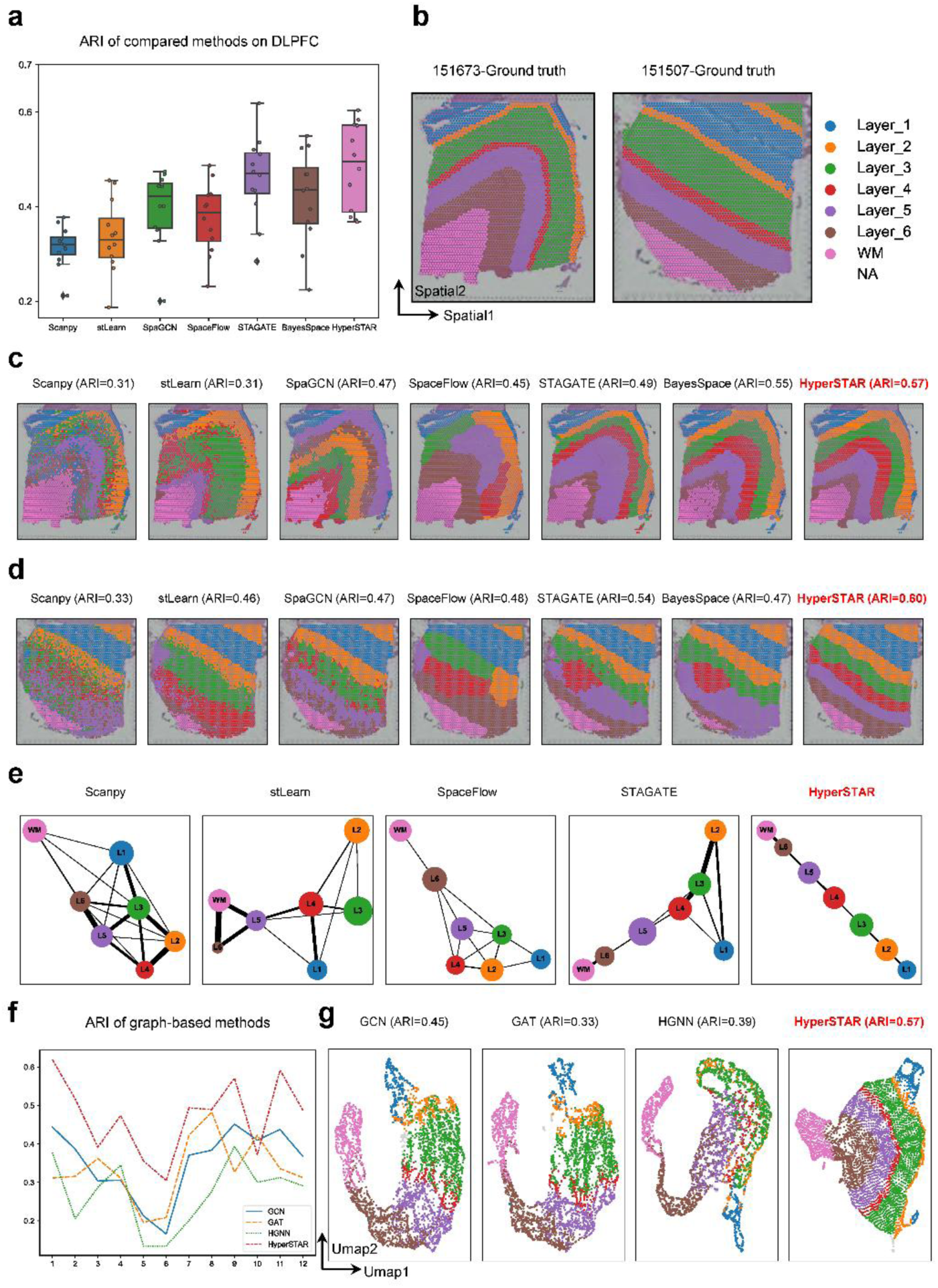
HyperSTAR achieves state-of-the-art performance on the DLPFC benchmark dataset. **a** Boxplots of clustering accuracy across all 12 sections of the dataset, measured by ARI scores for seven methods. **b** Ground-truth segmentation of cortical layers (from WM to Layer 1) and white matter in slices 151673 and 151507. **c** Spatial visualization of clustering results in slice 151673 compared to six state-of-the-art methods. **d** Spatial visualization of clustering results in slice 151507 compared to six state-of-the-art methods. **e** PAGA graphs generated by Scanpy, stLearn, SpaceFlow, STAGATE, and HyperSTAR. **f** Line plots of clustering accuracy across all 12 sections of the dataset, measured by ARI scores for four different graph-based networks (GCN, GAT, HGNN, and HyperSTAR). **g** UMAP plots of section 151673 obtained by the four graph-based networks.

We compared our method HyperSTAR with six commonly used packages for identifying spatial domains: Scanpy [23] (originally designed for single-cell omics), two methods that integrate histology images--stLearn and SpaGCN, two deep learning methods based on graphs--SpaceFlow and STAGATE, and one Bayesian statistical-based method, BayesSpace. HyperSTAR effectively identified the expected cortical layer structures, demonstrating significant improvement compared to other methods (Fig. 2a). For example, in sections 151673 and 151507 (ground truth shown in Fig. 2b), HyperSTAR clearly delineated the layer borders and achieved the highest clustering accuracy (ARI = 0.57 and ARI = 0.60, respectively) (Fig. 2c, 2d). The results of spatial visualization for all sections are provided in Supplementary Fig. S1-S15. We further confirmed the inferred trajectory using a trajectory inference algorithm named PAGA. The PAGA graphs illustrate that embeddings belonging to different clusters obtained by Scanpy, stLearn, and SpaceFlow are intermingled. As for STAGATE, it successfully separates WM, Layer 6, and Layer 5; however, Layer 1, Layer 2, Layer 3, and Layer 4 remain mixed. Our HyperSTAR embeddings effectively divide these cortical layers, providing a clear depiction of tissue topology (Fig. 2e).

We further conducted ablation experiments to illustrate the effectiveness of our strategies. In Fig. 2f, we present the performance of four graph-based networks: graph convolution networks (GCN) [30], graph attention networks (GAT) [31], hypergraph convolution networks (HGNN) [32], and our HyperSTAR, which combines hypergraph convolution operation and a graph attention mechanism. As shown in Fig. 2f, the ARI score of HyperSTAR outperforms other methods on almost all slices of the DLPFC dataset. In Fig. 2g, UMAP plots of embeddings obtained from four networks demonstrate the superiority of HyperSTAR. For GCN, GAT, and HGNN, there is some overlap between cortical layers, indicating some spots are mixed. In contrast, HyperSTAR demonstrates exceptional performance by clearly distinguishing all cortical layers without any overlap or mixing. To ensure the robustness of our findings, we have included multiple running results in Supplementary Fig. S20. Additionally, the denoising results and the detection of spatially variable genes (SVGs) are presented in Supplementary Fig. S21 and S22, respectively.

### HyperSTAR reveals sub-structure on finer resolutions of mouse olfactory bulb

To assess HyperSTAR’s performance across different resolutions, we selected a mouse olfactory bulb section from Stereo-seq, with finer spatial resolution at the single-cell level. Spatial domains on this section were initially identified using three methods: Scanpy, a widely used single-cell method; STAGATE, a graph-based deep learning method; and our proposed method, HyperSTAR. As depicted in Fig. 3b, Scanpy yielded discontinuous results, while STAGATE showed improved performance, albeit with some mixing between layers, such as the fusion of domain 3 and domain 4. In contrast, HyperSTAR identified a clear and continuous layer structure in the mouse olfactory bulb, corresponding to the mouse anatomy collected from the Allen Brain Atlas, as shown in Fig. 3a.

**Fig. 3.**
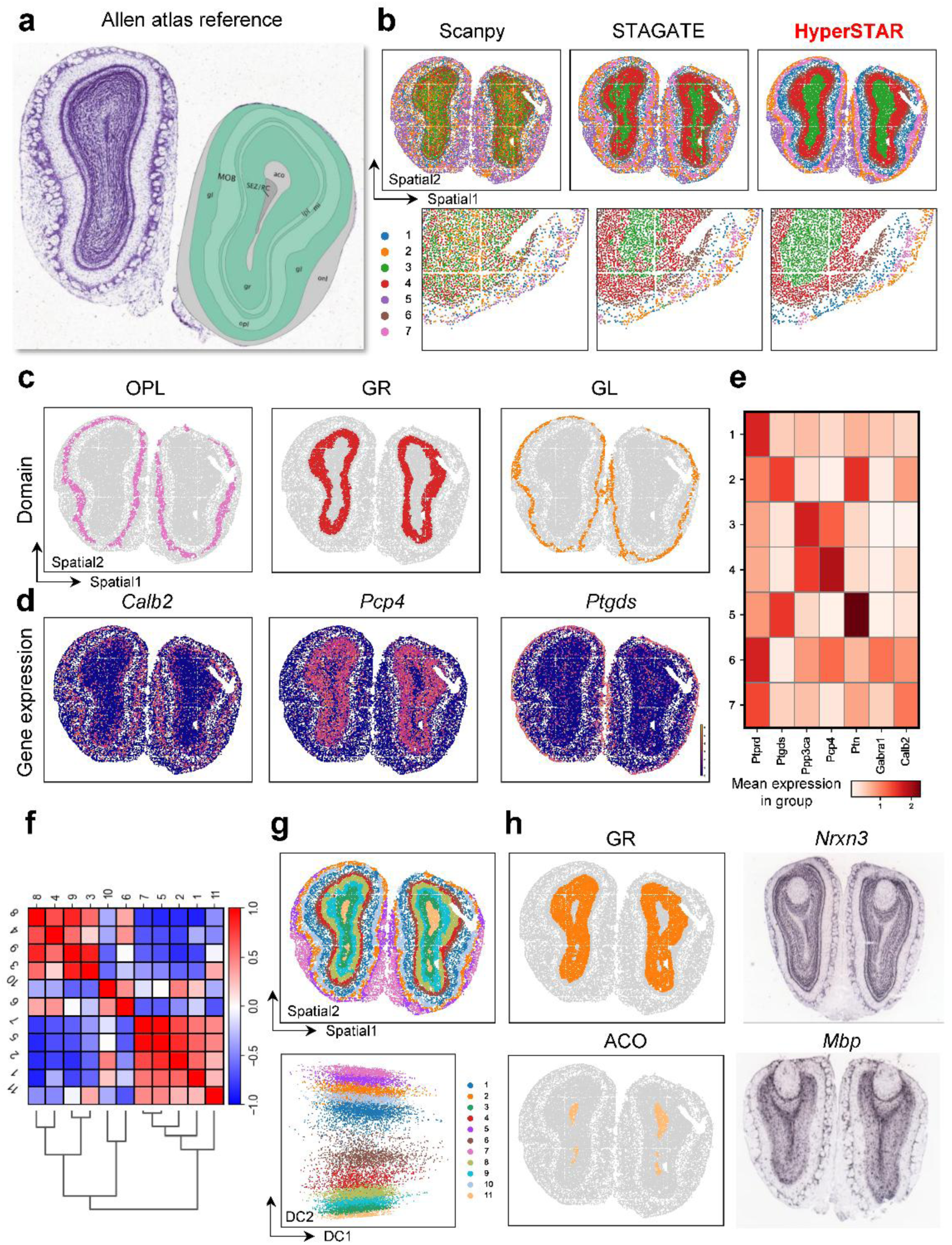
HyperSTAR reveals sub-structure ACO at finer resolutions of the Stereo-seq mouse olfactory bulb. **a** Mouse brain olfactory bulb anatomical structure from the Allen Brain Atlas. **b** Spatial visualization of results obtained by Scanpy, STAGATE, and HyperSTAR. **c** OPL, GL, and GR layers identified by HyperSTAR. **d** Spatial visualization of three marker genes expressed in OPL (Calb2), GL (Pcp4), and GR (Ptgds), respectively. **e** Heatmap displaying top genes characterizing seven groups identified by HyperSTAR. **f** Correlation matrix of HyperSTAR results, showing similarity within clusters and dissimilarity between clusters. **g** Top: spatial visualization results from HyperSTAR; Bottom: diffusion map plot revealing the sub-structure ACO within GR. **h** Top: GR identified by HyperSTAR and ISH image of marker gene Nrxn3 from the Allen Brain Atlas; Bottom: ACO identified by HyperSTAR and ISH image of marker gene Mbp from the Allen Brain Atlas.

To further analyze the results obtained by HyperSTAR, we ranked genes for characterizing groups, and the top genes identified for each group are shown in Fig. 3e. The gene expressions of *Calb2*, *Pcp4*, and *Ptgds*, shown in Fig. 3d, represent the three top genes identified by HyperSTAR for characterizing domains 7, 4, and 2, respectively. The distribution of domains 7, 4, and 2 identified by HyperSTAR is displayed in Fig. 3c, corresponding to the reference in Fig. 3a as layers OPL, GR, and GL. The spatial visualization and corresponding In Situ Hybridization (ISH) images from the Allen Atlas of marker genes, which distinguish the different layers, are also provided in Extended Data Figs. 1-2. These figures collectively illustrate HyperSTAR’s ability to reveal continuous tissue structure and find effective markers.

We adjusted the parameters for fine-grained clustering, and the result is shown in Fig. 3g. HyperSTAR reveals a sub-structure ACO within GR (Fig. 3h) and its top characterizing gene Mbp. Additionally, we identified the top gene in GR as *Nrxn3*. The ISH image illustrates the effectiveness of the result obtained by HyperSTAR. The heatmap of the correlation matrix of clustering also demonstrates the effectiveness of HyperSTAR, showing similarity within clusters and dissimilarity between clusters (Fig. 3f).

We further assessed HyperSTAR’s cross-resolution capability on various spatial resolutions of the above mouse olfactory bulb section. Across bin 20, bin 50, bin 100, and single-cell levels (Extended Data Figs. 3-4), our HyperSTAR results consistently exhibit a continuous structure aligned with the reference in Fig. 3a. Extending our evaluation to an adult axolotl brain slice further confirmed HyperSTAR’s efficacy, revealing a continuous structure corresponding to the reference (Extended Data Fig. 5).

### HyperSTAR unveils spatial intra-tumor heterogeneity in breast cancer section and its association with survival

To investigate the capability of HyperSTAR for tumor analysis, we applied it to a breast cancer section obtained from 10x Visium. This section was manually annotated by a pathologist into four distinct categories: ductal carcinoma in situ/lobular carcinoma in situ (DCIS/LCIS), invasive ductal carcinoma (IDC), healthy tissue, and the tumor edge (Fig. 4a) [33]. Remarkably, HyperSTAR accurately identified these categories, demonstrating strong consistency with the manual annotations (Fig. 4b and Supplementary Fig. S23).

**Fig. 4.**
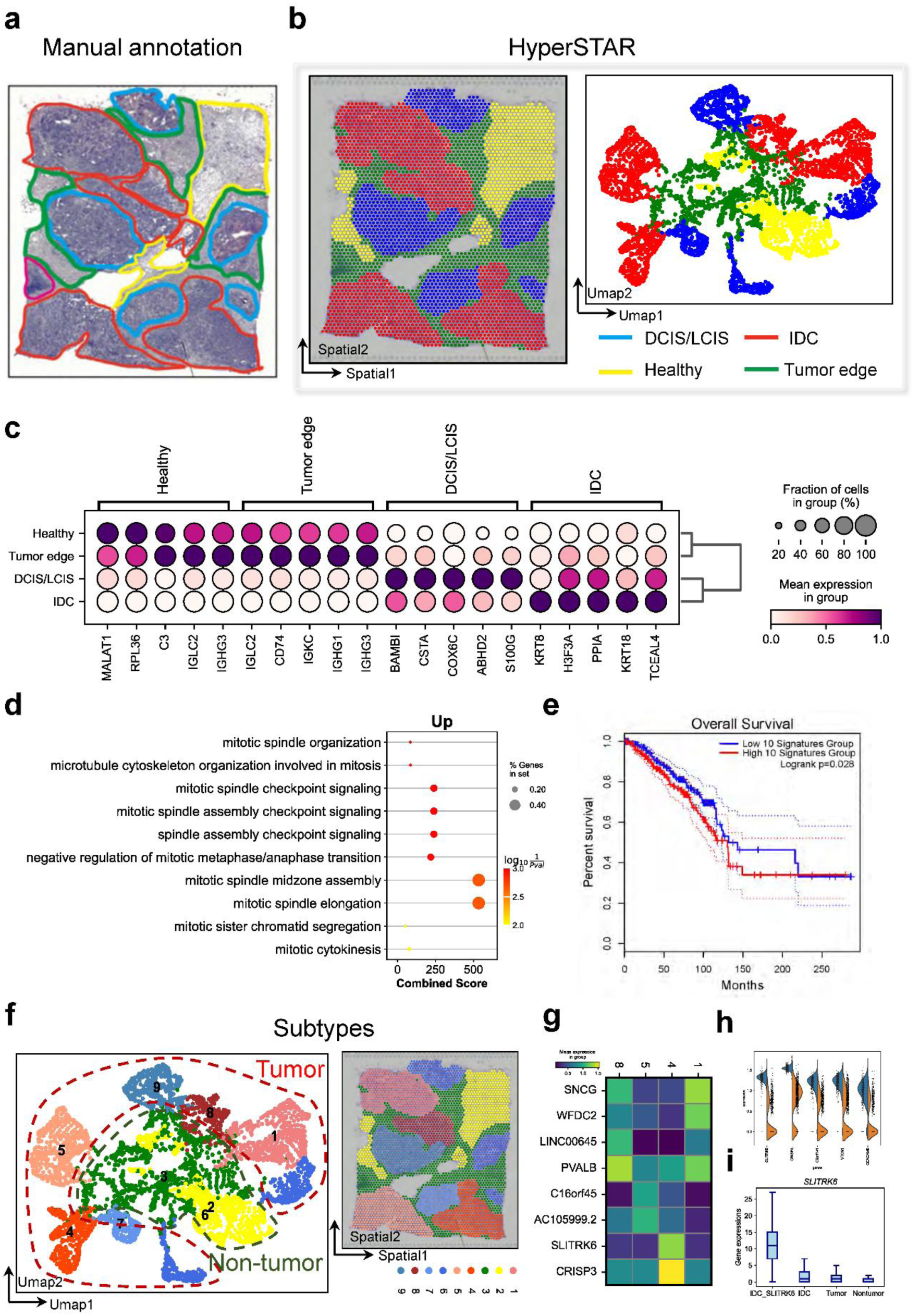
HyerSTAR identifies gene markers in breast cancer subtypes and their association with survival. **a** Manual annotation of breast cancer section from 10x Visium by pathology experts. **b** Spatial results and UMAP plots generated by HyperSTAR for this section. **c** Dot plots of marker genes identified by HyperSTAR. **d** Gene ontology enrichment analysis of the differentially expressed genes (DEGs) between IDC and the rest. **e** Overall survival rates of patients with 10 signature genes for IDC, analyzed using TCGA breast cancer gene expression data via GEPIA2. **f** Subtypes identified by HyperSTAR in the breast cancer section. **g** Heatmaps of marker genes in IDC subtypes. **h** Top five marker genes of subtype IDC_SLITRK6 compared to the rest of IDC. **i** Gene expression levels of SLITRK6 in IDC_SLITRK6, IDC, tumor, and non-tumor regions.

We next interrogate the diagnosis power of HyperSTAR’s results. We rank and plot top 5 genes for characterizing these 4 domains (Fig. 4c). For *KRT8* and *KRT18*, which are found highly expressed in IDC region. *KRT8* is an independent prognostic indicator for poor survival in lung adenocarcinoma tissue [34]. *KRT18* is often together with *KRT8*, involved in interleukin-6 (IL-6)-mediated barrier protection. This patient has been clinically diagnosed as Grade III. Logan and et al. claims that *KRT8/18* expression differentially distinct subtypes of grade III invasive ductal carcinoma of the breast cancer [35], which is consistent with our results. We further compared transcriptional differences between IDC and the rest by performing differential expression analysis followed by pathway enrichment analysis. The differential expression genes (DEGs) of IDC are enriched in the process of mitosis, indicating abnormal proliferation of cancer cells (Fig. 4d), which indicates the area is probably infiltrated with cancer cells and also highly active. We further investigate the correlation of these DEGs with survival on an independent clinical data from TCGA breast cancer cohort. 10 signature genes (Supplementary Table 1) were selected to cutoff patients, and the result shows that low expression of these 10 signatures correlated with shorter overall survival (Fig. 4e). Hence, these results indicate that there is possible cell development from cancer stem cells to malignant cells. Here are also some research claims that *KRT8* is a pan-cancer indicator [36], In light of this, we conducted a pan-cancer survival analysis using these DEGs identified in IDC. The outcomes further reveal a correlation between these DEGs and pan-cancer development (Suppementary Fig. S16).

We further investigate the subtypes of IDC region. From our HyperSTAR, the IDC is divided into 4 subtypes (Fig. 4f). The heatmap, generated by the top two genes of each subtype (Figure 4g), highlights the unique expression profiles within each subtype. Notably, for Domain 4, *SLTRK6* and *CRISP3* exhibit high expression levels, with *SLITRK6* emerging as a potential marker for this specific subtype. The accompanying boxplot illustrates that *SLTRK6* is significantly upregulated in IDC_SLTRK6 compared to IDC, tumor, and non-tumor regions (Figure 4i). In summary, HyperSTAR not only enables the identification of intra-tumor heterogeneity but also facilitates the exploration of distinct subtypes within the intra-tumoral environment.

### HyperSTAR is generalisable

To comprehensively validate the versatility of HyperSTAR across various technologies and different omics, we applied it to multiple platforms, including 10x Genomics (10x Visium, 10x Xenium), MERFISH [37], NanoString (CosMx), and osmFISH (Supplementary Fig. S17, S23, and Fig. 5). Our evaluation further extended to different omics layers, encompassing both transcriptomics and proteomics, demonstrating the robust adaptability and broad applicability of HyperSTAR.

**Fig. 5.**
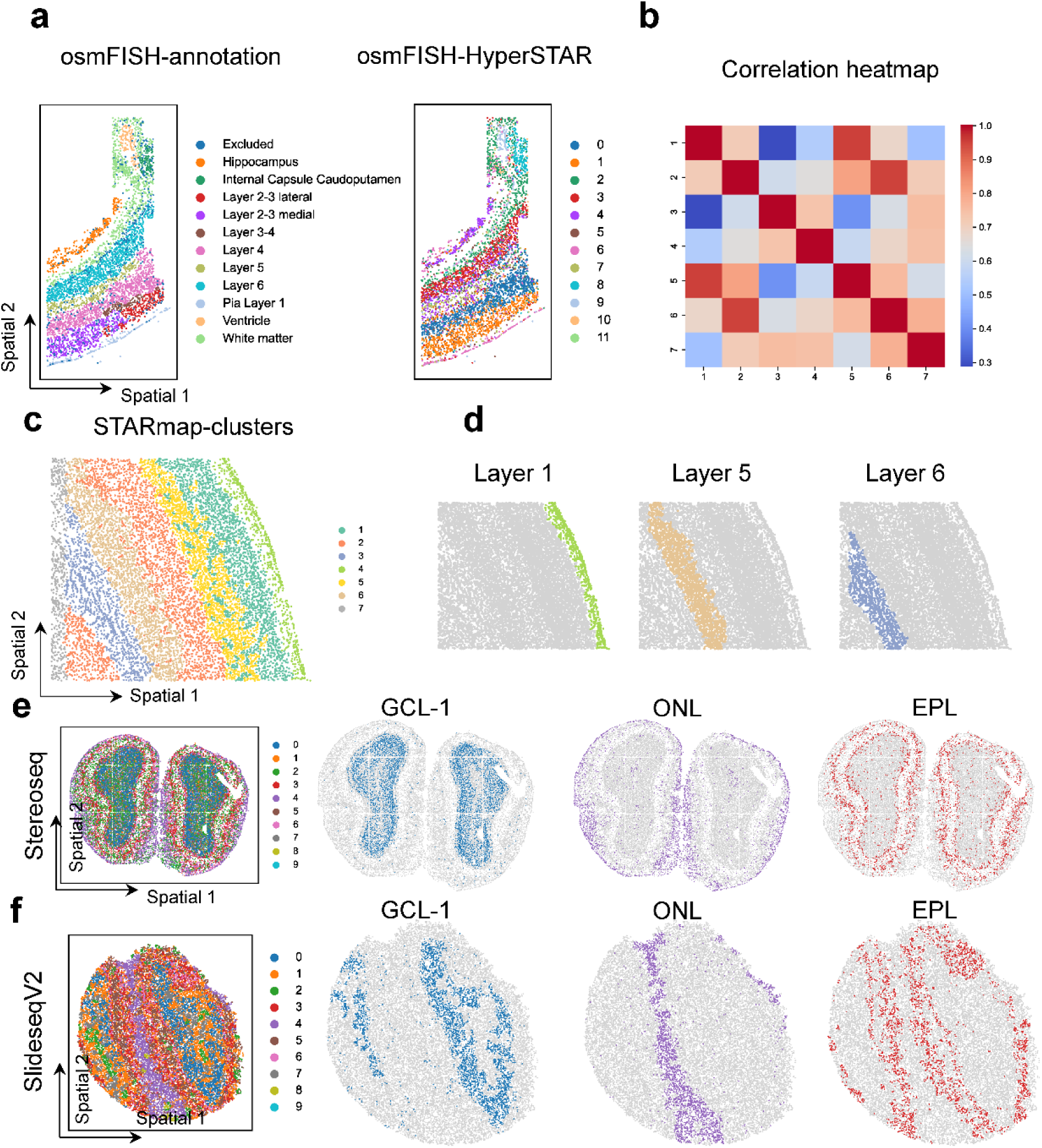
HyperSTAR is applicable to various spatial omics datasets and demonstrates joint clustering capabilities. **a** Manual annotation and HyperSTAR identification of the somatosensory cortex from osmFISH data. **b** Correlation heatmaps of clustering results obtained by HyperSTAR on STARmap data. **c** Spatial results of HyperSTAR on a mouse brain section from STARmap. **d** Spatial distribution of different layers revealed by HyperSTAR on STARmap data. e Joint clustering results of Stereo-seq data and spatial distribution of different layers identified by HyperSTAR. **f** Joint clustering results of SlideSeqV2 data and spatial distribution of different layers identified by HyperSTAR.

HyperSTAR reveals the cortex distribution of the somatosensory cortex (Fig. 5a) in a dataset produced by osmFISH, a technique based on multiplexed fluorescence *in situ* hybridization [38]. osmFISH enables the detection of hundreds of genes at single-cell resolution, providing valuable insights into the spatial organization of complex tissues. Applying HyperSTAR to this dataset, we were able to identify distinct layers and regions within the somatosensory cortex, showcasing the method’s ability to uncover biologically meaningful structures. To further demonstrate the versatility of HyperSTAR, we applied it to the STARmap [39] mouse brain data. As a result, HyperSTAR explicitly revealed the distribution of brain layers (Fig. 5c, 5d), highlighting its effectiveness in identifying spatial patterns across different experimental platforms. The heatmaps of the result in STARmap data display the correlation matrix of the identified clusters, with high correlation values within clusters but low correlation values between clusters (Fig. 5b). This indicates that HyperSTAR successfully captures the underlying structure of the data, grouping together domains with similar spatial gene expression profiles while separating domains from different layers or regions.

The Hypergraph framework is a highly scalable and versatile approach that be applied to jointly cluster data from multiple sources. To evaluate its capability, we selected mouse olfactory bulb SRT data from two different sequencing platforms: SlideSeqV2 and Stereo-seq. These datasets exhibit differences in quality and distribution (Supplementary Fig. S18). To integrate data from these different sequencing platforms, we extended the hypergraph construction process beyond single slices. In addition to creating hyperedges within each slice, we established inter-slice relationships by constructing hyperedges that connect spots across slices based on their normalized log-transformed gene expression profiles. This approach allows for the identification of shared spatial patterns across different datasets.

To address potential batch effects arising from the different sequencing platforms, we utilized Harmony for batch effect correction. The joint clustering results generated by HyperSTAR (Fig. 5e, 5f, and Supplementary Fig. S24) successfully delineated distinct layers within the mouse olfactory bulb, aligning with its known anatomical structure. By leveraging the complementary information provided by SlideSeqV2 and Stereo-seq, HyperSTAR effectively identifies shared spatial patterns across datasets, offering a more comprehensive understanding of tissue organization.

## Discussion

The advent of spatial omics technologies has revolutionized our understanding of biological processes by enabling the simultaneous acquisition of biomolecular and spatial information. Accurate identification of spatial domains is crucial for deciphering spatiotemporal life processes. In this study, we introduce HyperSTAR, a powerful and versatile method for spatial omics analysis that excels in accurately identifying spatial domains, even in high-resolution spatial data. By incorporating a gene expression-guided hyperedge decomposition module and a hypergraph attention convolutional neural network, HyperSTAR demonstrates exceptional precision in recognizing spatial domains across varying resolutions. The advancements hold significant implications for exploring organ development and tumor microenvironments.

Benchmarking and comparisons with existing advanced graph neural network models underscore HyperSTAR’s superior performance in uncovering tissue substructures, inferring spatiotemporal patterns, and denoising spatially resolved gene expressions. Its application to spatial omics data from diverse platforms—including 10x Genomics, NanoString, CODEX, and osmFISH—highlights its adaptability and robustness across different technologies. Furthermore, the validation of HyperSTAR’s findings through functional and survival analyses of independent clinical data, particularly in the context of breast cancer, reinforces its utility in deciphering spatial heterogeneity and its potential for clinical relevance.

A key strength of HyperSTAR lies in its ability to capture intricate higher-order relationships among spatially neighboring multi-spots. However, the current version of HyperSTAR is primarily focused on identifying spatial domains and has yet to fully exploit the potential of hypergraphs. In future work, we aim to extend HyperSTAR to integrate multi-omics data from spatial omics experiments generated by different platform technologies. Additionally, incorporating histology images could provide valuable insights for addressing clinically relevant questions and further enhancing the method’s applicability.

In conclusion, HyperSTAR represents a significant advancement in spatial omics analysis, addressing challenges associated with diverse spatial resolutions and enabling the exploration of complex biological systems. Its ability to capture intricate higher-order relationships within spatially neighboring multi-spots makes it an invaluable tool for researchers studying organ development and tumor microenvironments, and other spatially resolved biological processes. This study establishes HyperSTAR as a versatile and effective computational approach with broad applicability across diverse spatial omics datasets and large-scale analyses. Future directions include refining and expanding HyperSTAR’s capabilities, integrating multi-omics data and histology images, and exploring its application in additional biological contexts. These efforts will further solidify HyperSTAR’s role as a transformative tool in spatial omics research.

## Acknowledgements

This work has been supported by the National Key Research and Development Program of China (2022ZD0117700), the National Natural Science Foundation of China (U21A20520, 62325204), the Key Area Research and Development Program of Guangzhou City (202206030009, 2023B01J1001), Australian National Health and Medical Research Council (NHMRC) Ideas grant APP 2020646, Australian Rsearch Council Linkage project grant (LP220200614), Major and Seed Inter-Disciplinary Research Projects awarded by Monash University.

## Author contributions

Hongmin Cai, Zhiyuan Yuan, and Jiangning Song jointly supervised the research and provided guidance throughout the study. Yi Liao, Fei Qi, and Weitian Huang conceptualized and designed the research. Yi Liao and Chong Zhang performed the research, including data collection, analysis, and interpretation. Zhikang Wang, Jinyu Li, and Shangyan Cai made significant contributions to image enhancement and manuscript language refinement. Yi Liao wrote the initial draft of the manuscript, and all authors critically reviewed and revised the manuscript for important intellectual content.

## Supplementary information

Supplementary Figs. S1-S24 and Table S1.

## Source data

Source Data Fig. 3-4, related to Figures 3 and 4.

**Extended Data Fig. 1.**
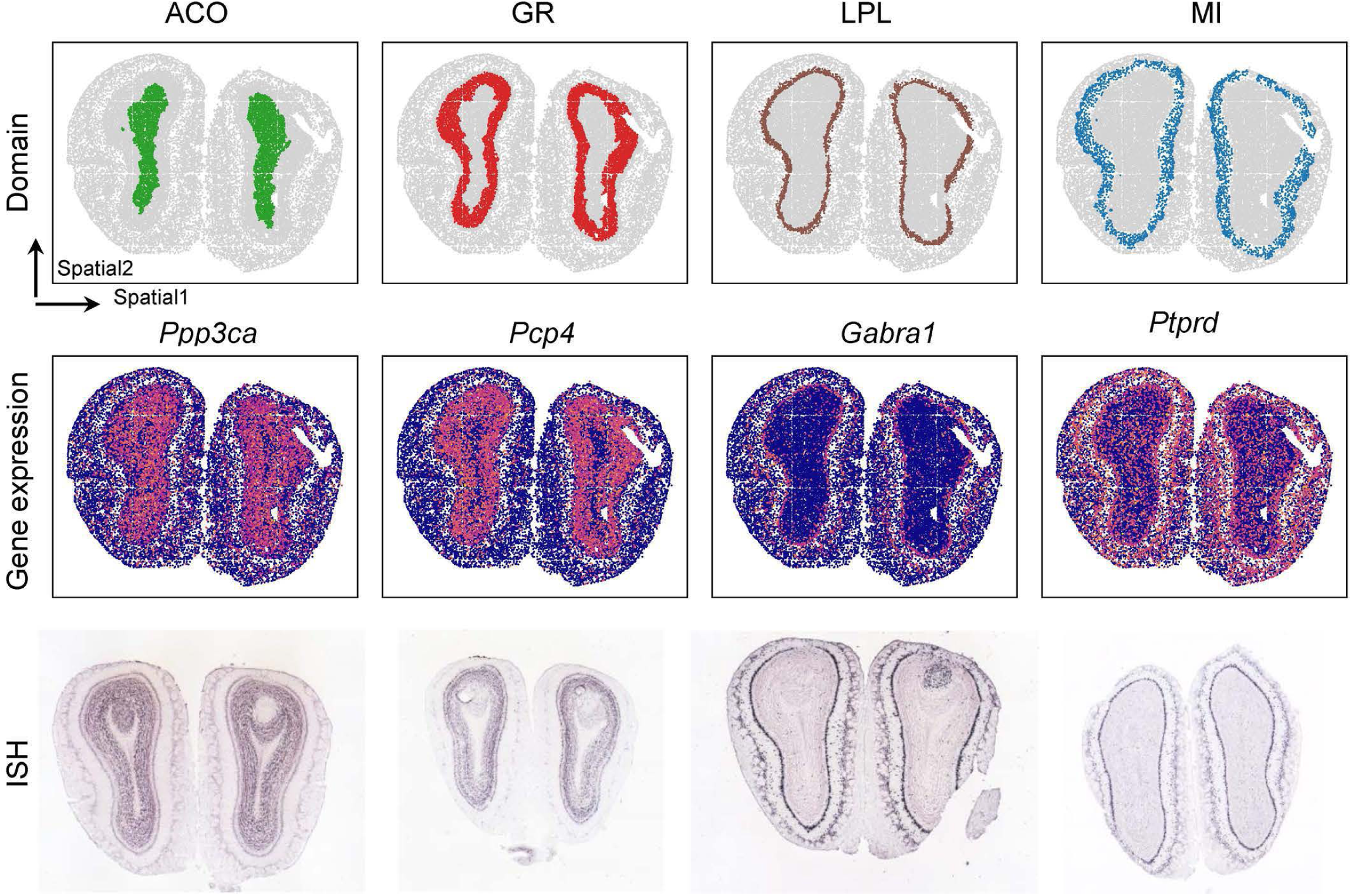
HyperSTAR reveals different layers at finer resolutions of the Stereo-seq mouse olfactory bulb. Spatial visualization of ACO, GR, LPL, and MI identified by HyperSTAR.

**Extended Data Fig. 2.**
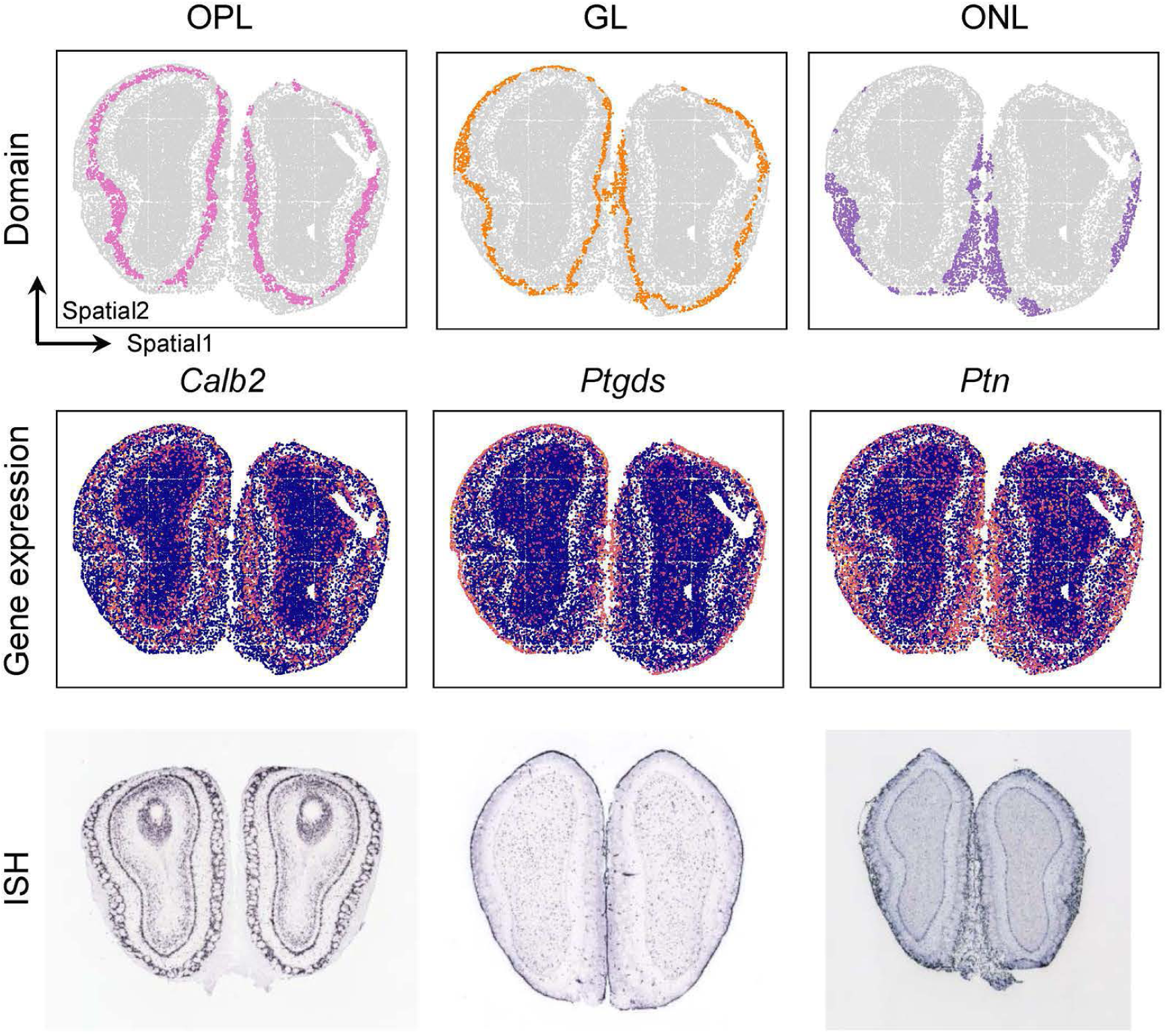
HyperSTAR reveals different layers at finer resolutions of the Stereo-seq mouse olfactory bulb. Spatial visualization of OPL, GL, and ONL identified by HyperSTAR.

**Extended Data Fig. 3.**
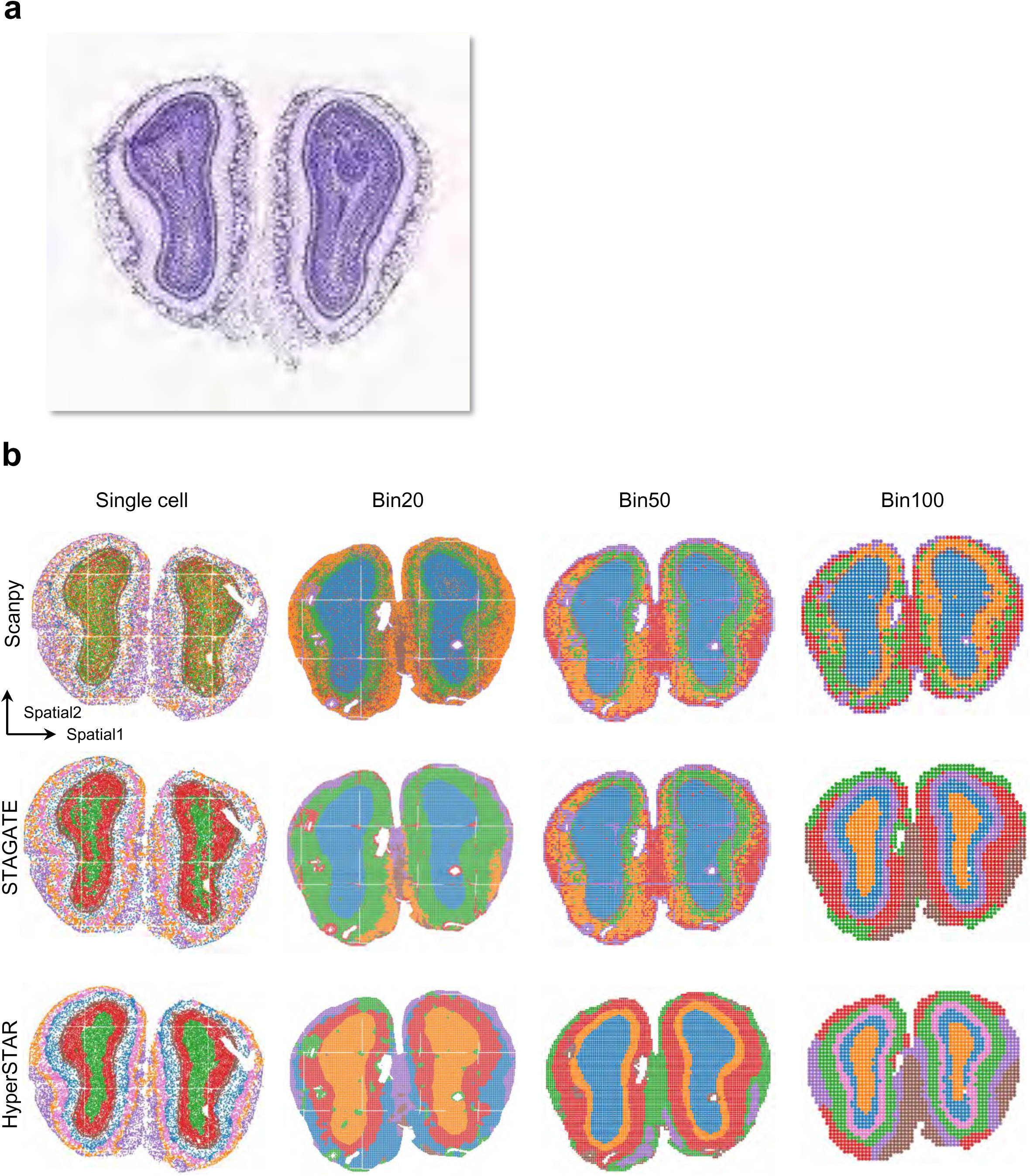
HyperSTAR achieves robust performance across different resolutions on Stereo-seq data. **a** Reference from the Allen Brain Atlas. **b** Spatial visualization of results obtained by Scanpy, STAGATE, and HyperSTAR across four different resolutions (Single cell, bin 20, bin 50, and bin 100).

**Extended Data Fig. 4.**
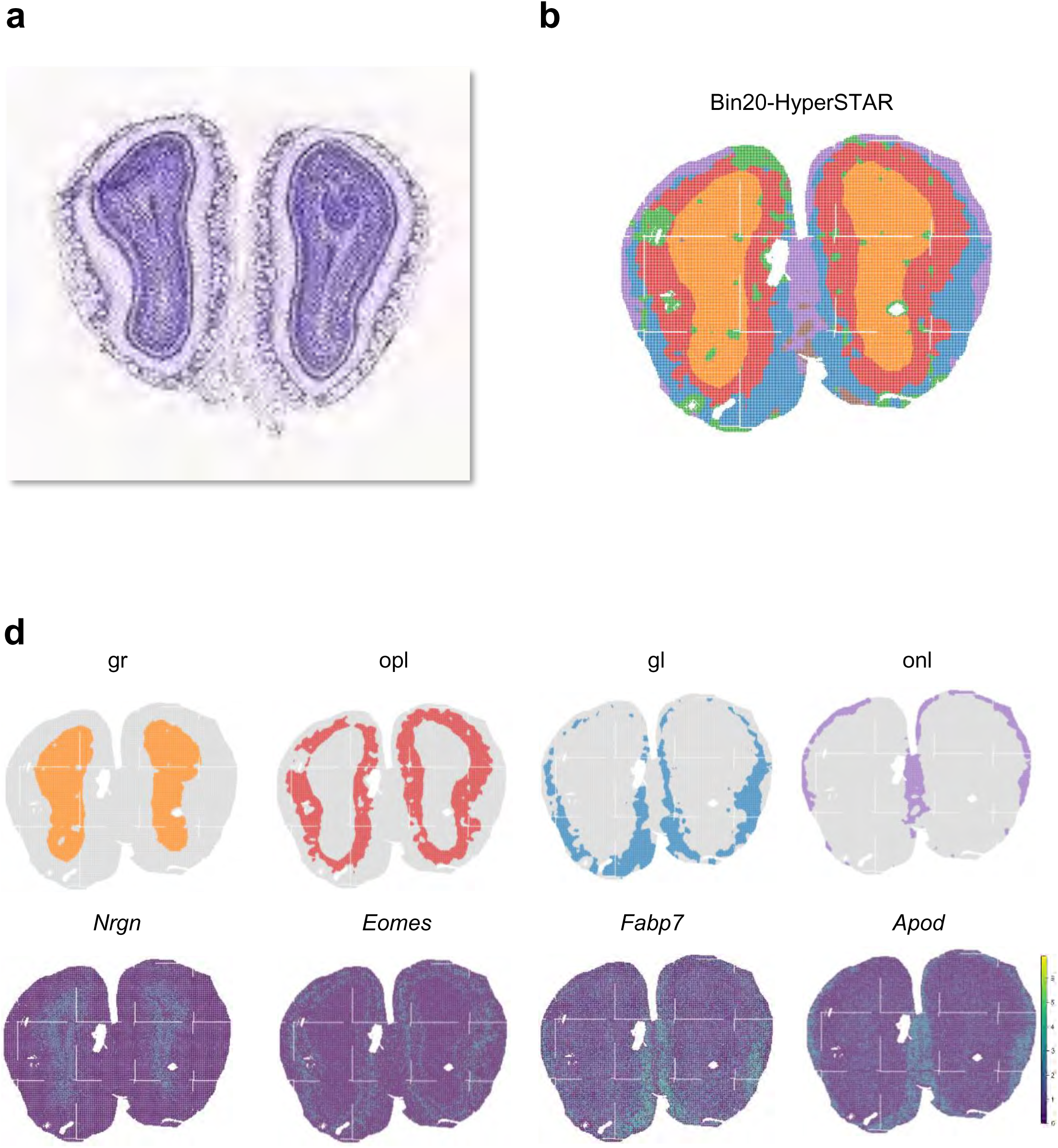
HyperSTAR achieves roubust performance across different resolutions on Stereo-seq data. **a** Reference from the Allen Brain Atlas. **b** Results obtained by HyperSTAR on bin 20 mouse olfactory bulb data. **c** Spatial visualization of different layers identified by HyperSTAR and their corresponding marker gene expression.

**Extended Data Fig. 5.**
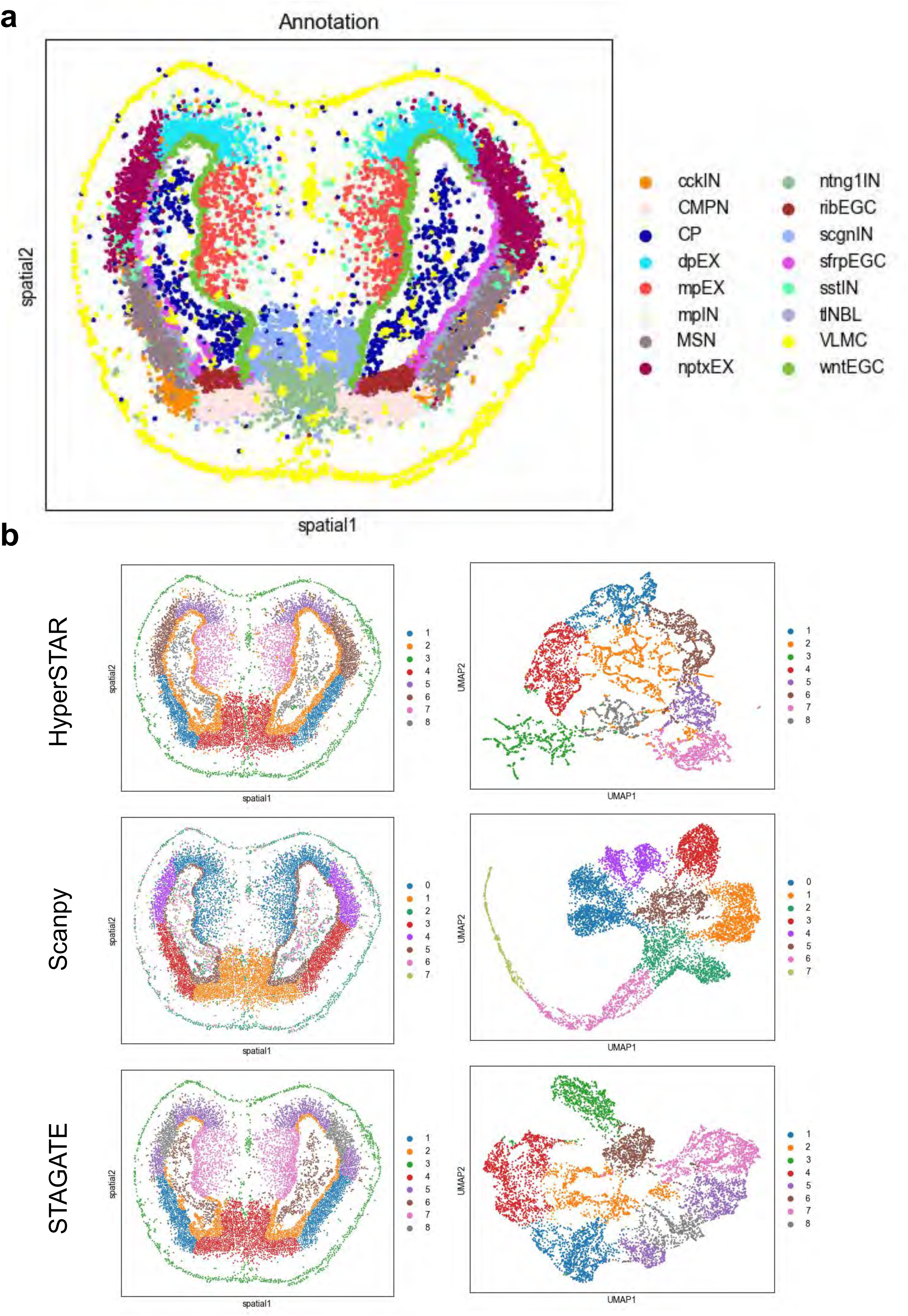
HyperSTAR reveals layers of adult axolotl brain. **a** Annotation of adult axolotl brain data. **b** Spatial visualization of results obtained by Scanpy, STAGATE, and HyperSTAR.

## STAR Methods

### Data description

We applied HyperSTAR to a diverse range of spatial omics datasets generated by various technologies to demonstrate its effectiveness. These datasets span multiple platforms, including SRT technologies such as 10x Visium, Stereo-seq, osmFISH, MERFISH, and Xenium, as well as CODEX, which is used for spatial proteomics. These platforms produce data with varying resolutions, ranging from spot-level to single-cell resolution. For example, 10x Visium datasets have a resolution of 55 µm (1-10 cells per spot), while Stereo-seq provides spot resolutions with bin sizes of 20, 50, 100, and single-cell resolution.

Specifically, we used the human dorsolateral prefrontal cortex (DLPFC) dataset [29], generated by 10x Visium, as the benchmark for spatial domain identification. This dataset includes 12 human DLPFC sections from three individuals, with manual annotations provided by the original authors. The 10x Visium mouse brain coronal dataset comprises 2,702 spots and 21,949 genes as raw features. The Stereo-seq mouse brain olfactory bulb dataset, with single-cell resolution, includes 19,109 spots and 27,106 genes as raw features. The breast cancer section dataset, generated by 10x Visium, has a median of 20,762 unique molecular identifiers (UMI) counts per spot and a median of 6,026 genes per spot. These datasets cover diverse species, organs, and resolutions, showing the versatility of HyperSTAR.

### Data preprocessing

To effectively analyze the raw data, we employed a series of preprocessing steps. First, we removed spots located outside the primary tissue region to focus on biologically relevant areas. Next, we performed feature selection to identify the top highly variable genes (HVGs) (typically 3,000 genes). After selecting the HVGs, we applied a log transformation to the gene expression values. The resulting normalized HVGs were then used as input node representations in the hypergraph learning module. This preprocessing ensures that the hypergraph learning module can effectively capture the intricate relationships among spots and identify biologically meaningful spatial domains.

### Hypergraph construction

After preprocessing to obtain spot representations, we constructed the hypergraph structure in two steps:

1. Initial Hyperedge Construction: For each spot, we generated a hyperedge using the k-nearest neighbors (kNN) approach based on spatial information of the spots. This step captures local spatial relationships among spots, ensuring that spatially adjacent spots are connected in the hypergraph.
2. Hyperedge Refinement: To address the scenarios where spatially adjacent spots belong to different biological domains, we refined the hyperedge set using pre-clustering results based on gene expression. This refinement aligns the hypergraph structure with underlying biological domains, thereby improving the accuracy of the spatial domain identification. Specifically:

If the spots within a hyperedge belong to the same cluster, the hyperedge remains unchanged, indicating that spatially adjacent spots share similar gene expression patterns and likely belong to the same biological domain.

If the spots within a hyperedge belong to different clusters, we consider two scenarios:

i. Single Outlier Spot: If only one spot belongs to a different cluster, it is removed from the hyperedge to maintain consistency. This ensures that the hyperedge only contains spots that are both spatially adjacent and share similar gene expression patterns;
ii. Multiple Outlier Spots: If multiple spots belong to different clusters, we decomposed into two new hyperedges, each containing from the same cluster. This effectively separates the spots that are spatially adjacent but have distinct gene expression patterns. By refining the hyperedge set based on the pre-clustering results, we better capture the spatial and biological relationships among spots, leading to a more accurate representation of spatial domains in the hypergraph structure.

### Hypergraph learning

Once the hypergraph is constructed, we input the spot features and hypergraph structure into the hypergraph learning module, which consists of two parts: the encoder and the decoder. The encoder generates latent low-dimensional embeddings of spots used for identifying spatial domains and downstream analyses. The decoder produces reconstructed features of spots utilized for denoising.

Given a hypergraph **G** = (**X**, **H**), where 𝐆 ∈ 𝐑^𝑛×𝑑^ represents the feature matrix of *n* spots and

𝐇 ∈ 𝐑^𝑛×𝑚^ represents incidence matrix from vertexes to hyperedges defined as [32, 40]:

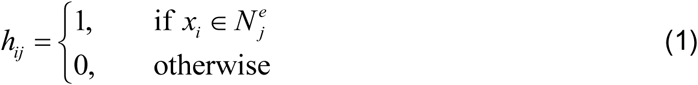

where *N* denotes the set of vertexes connected to the *j*-th hyperedge.

#### Encoder

The encoder takes the normalized gene expressions 𝐗_0_ as input and generates the spot embeddings by collectively aggregating the information from its neighbors in the same hyperedge. To adaptively learn the weights of the hyperedges, we used three layers of hypergraph attentional layers.

#### Hypergraph convolutional layer

Message propagation on the hypergraph is more complex than the graph, involving two stages: attentive hyperedge aggregation and attentive vertex aggregation [41]. In the attentive hyperedge aggregation, the connected vertexes in the same hyperedge are aggregated to generate the hyperedge embeddings. As for the *t*-th layer, firstly, the hyperedge features **E** can be calculated by

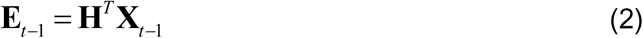

where 𝐗_𝑡−1_ represents output vertex features of the last layer. For each pair of (𝑒_𝑖_, 𝑒_𝑗_) (𝑒_𝑗_ ∈

𝑁^𝑣^), we calculate the coefficient 𝑎_𝑖𝑗_ between the hyperedge 𝑖 and hyperedge 𝑗 by introducing the attention mechanism [42]:

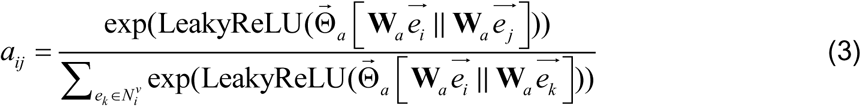

where 𝐖 ∈ 𝐑^𝑑′×𝑑^ is a shared linear transformation, parametrized by a weight matrix. 𝜃⃗ ∈

𝑅^2𝑑′^ represents a shared attentional mechanism, parametrized by a vector. 𝑁_i_^𝑣^ denotes the set of hyperedges connected to the *i*-th vertex. The hyperedge features aggregated by attentive hyperedge aggregation can be calculated as follows:

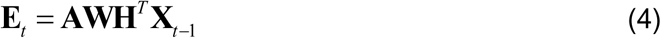

The attentive vertex aggregation module aggregates the connected hyperedge information to generate the vertex embeddings. First, the vertex features can be calculated by

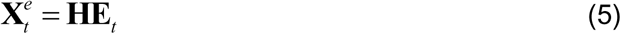

We calculate the coefficient 𝑏_𝑖𝑗_ between the hyperedge 𝑖 and vertex 𝑗 based on the attention mechanism:

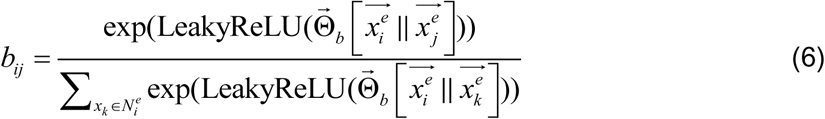

where 𝜃⃗_𝑏_ ∈ 𝑅^2𝑑^ represents a shared attentional mechanism, parametrized by a vector, while

𝑁_i_^𝑒^ denotes the set of vertexes connected to the i-th hyperedge. Next, the output vertexes features 𝐗_𝑒_ can be calculated by

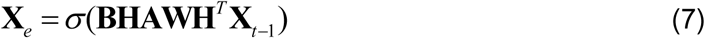

where σ is the activation function.

#### Decoder

The decoder takes the low-dimensional latent embeddings of spots as input and reconstructs the spot features in the original input dimensions of the encoder. We also used hypergraph attentional networks as input to build a symmetrical decoder for encoder. After applying three layers of hypergraph attentional networks, the output of the last layer is considered as the reconstructed spot features. As for the output of t-th layer in the decoder, it could be obtained by:

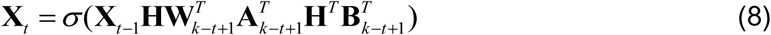

where **X**_0_ is the latent embeddinngs learned by the encoder, while k represents the number of layers of the encoder or decoder in our model. We set k = 3 in this study.

#### Loss function

Hypergraph structured data include node features and the graph structure, both of which should be encoded by high-quality node representations. We minimized the reconstruction loss of node features as follows:

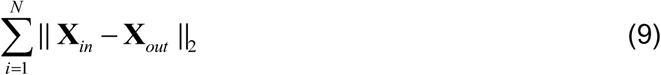

where 𝐗_𝑖𝑛_ is the input normalized gene expression 𝐗_0_ of the encoder, and represents the final output of the decoder.

### Downstream analysis

#### Clustering

To identify spatial domains based on the embeddings obtained by HyperSTAR, we employed different strategies depending on the availability of prior information about the number of labels or domains. When the number of labels is known, we utilized the mclust clustering algorithm [43]. For data without prior information about the number of domains, we used the Louvain algorithm implemented in the SCANPY package. By employing these two different clustering strategies based on the availability of prior information, we can flexibly adapt the spatial domain identification process to various datasets and research questions.

#### Spatial trajectory inference

To visualize the spatial trajectory inference, we employed two complementary approaches: the UMAP [44] plot and PAGA graph [45]. The UMAP plot is used to visualize the embeddings generated by the HyperSTAR. The PAGA graph is another powerful tool for visualizing spatial trajectories. PAGA is a graph-based method that combines clustering with trajectory inference, allowing for the identification of both discrete cell states and continuous transitions between them. In the context of spatial omics data, the PAGA graph can reveal the relationships between spatial domains and potential trajectories connecting them. We generated the PAGA graph using the scanpy.pl.paga function from the Scanpy package, which takes the HyperSTAR embeddings as input and produces a visually interpretable graph representation of the spatial trajectories. By combining the UMAP plot and PAGA graph, we can gain complementary and valuable insights into the spatial organization and potential trajectories present in the data, thereby facilitating the interpretation of the underlying biological processes.

#### SVG detection

We performed the Wilcoxon test implemented in the Scanpy package to identify spatially variable genes for spatial domains with a 1% FDR threshold (Benjamin-Hochberg adjustment).

#### Denoising

The denoised gene expression is generated by the decoder, and the visualization demonstrates that the denoised features are more effective and representative in distinguishing different domains.

## Data availability

All datasets analyzed in this work are publicly available in raw form from their original studies. Specifically, the 10x Visium human DLPFC dataset [29] is accessible within the *spatialLIBD* package (http://spatial.libd.org/spatialLIBD). The Slide-seqV2 mouse brain dataset \cite is obtained from the Broad Institue Single Cell Portal (https://singlecell.broadinstitute.org/single\_cell/study/SCP815/sensitive-spatial-genome-wide-expression-profiling-at-cellular-resolution\#study-summary). The Stereo-seq mouse brain dataset is collected from https://github.com/JinmiaoChenLab/SEDR\_analyses. The Stereo-seq axolotl data is obtained from https://db.cngb.org/stomics/artista/. The osmFISH mouse brain data is collected from https://linnarssonlab.org/osmFISH/. The Xenium mouse brain data is collected from https://www.10xgenomics.com/datasets/fresh-frozen-mouse-brain-replicates-1-standard. The CODEX Barrett’s esophagus data [46] is obtatined from https://datadryad.org/stash/share/1OQtxew0Unh3iAdP-ELew-ctwuPTBz6Oy8uuyxqliZk. Other data is from SODB [47]. The mouse brain reference is available in the Allen brain atlas (https://human.brain-map.org). The breast tumor section dataset is available on 10x Genomics weibsite https://www.10xgenomics.com/resources/datasets/human-breast-cancer-block-a-section-1-1-standard-1-1-0 and The annotation of breast tumor section by experts is obtained by DeepST [33] (https://academic.oup.com/nar/article/50/22/e131/6761985).

## Code availability

The HyperSTAR method is implemented in Python and is available on Github: https://github.com/Ringoio/HyperSTAR.

## Acknowledgements

This work has been supported by the National Key Research and Development Program of China (2024YFF1206600), the National Natural Science Foundation of China (U21A20520, 62325204), the Key Area Research and Development Program of Guangzhou City (2023B01J1001), Australian National Health and Medical Research Council (NHMRC) Ideas grant APP 2020646, Australian Rsearch Council Linkage project grant (LP220200614), Major and Seed Inter-Disciplinary Research Projects awarded by Monash University.

## Extended data

Extended Data Fig. 1-5.

## Supplementary information

Supplementary Figs. S1-24 and Table 1.

## Source data

Source Data Fig. 3-4.

