## Supplementary figures for "Unveiling Tissue Structure and Tumor Microenvironment from Spatial Omics by Hypergraph Learning": Supplementary figures.pdf

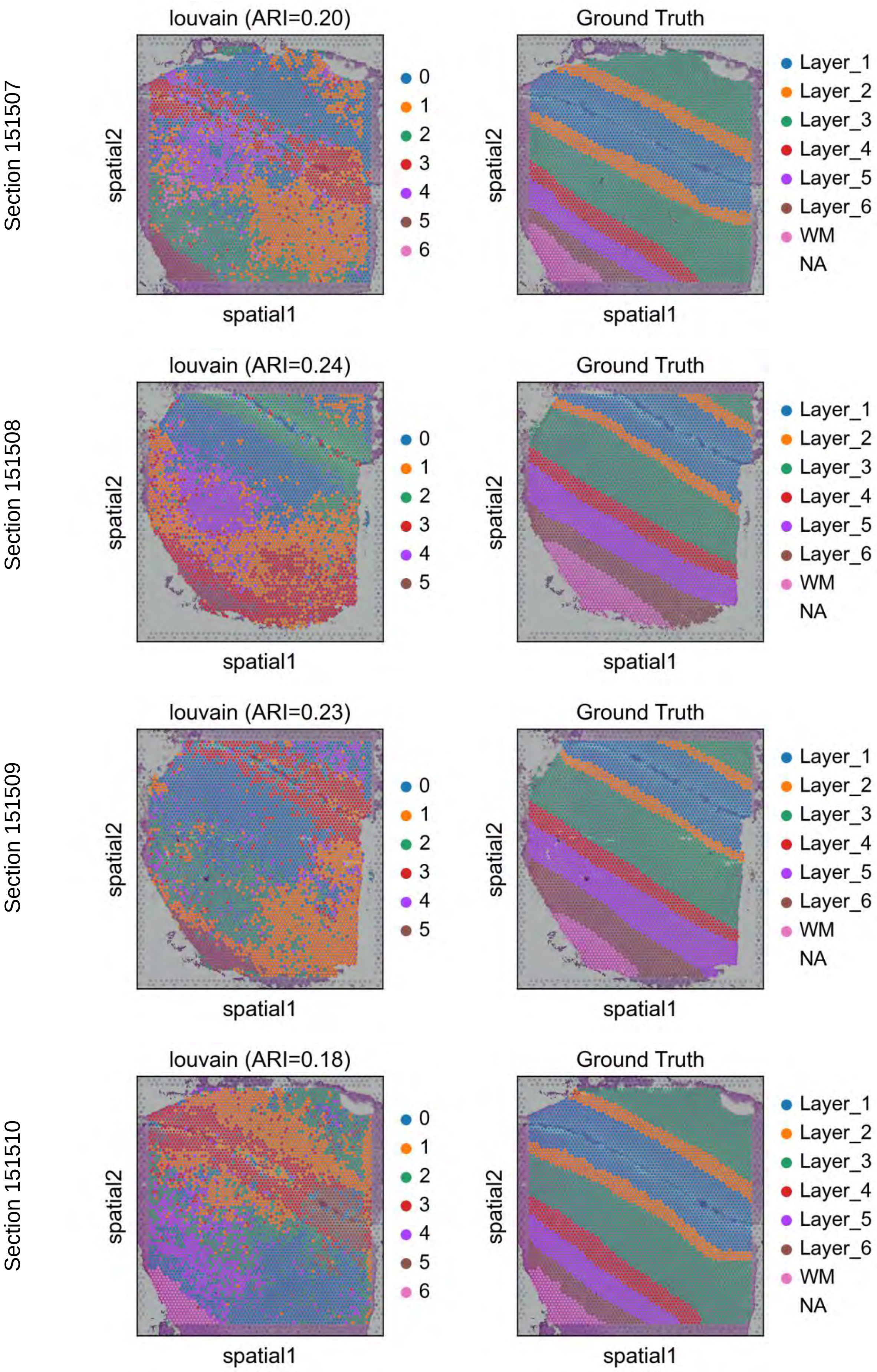


Supplementary Fig. S1. The result of Scanpy on DLPFC dataset sections 151507-151510.

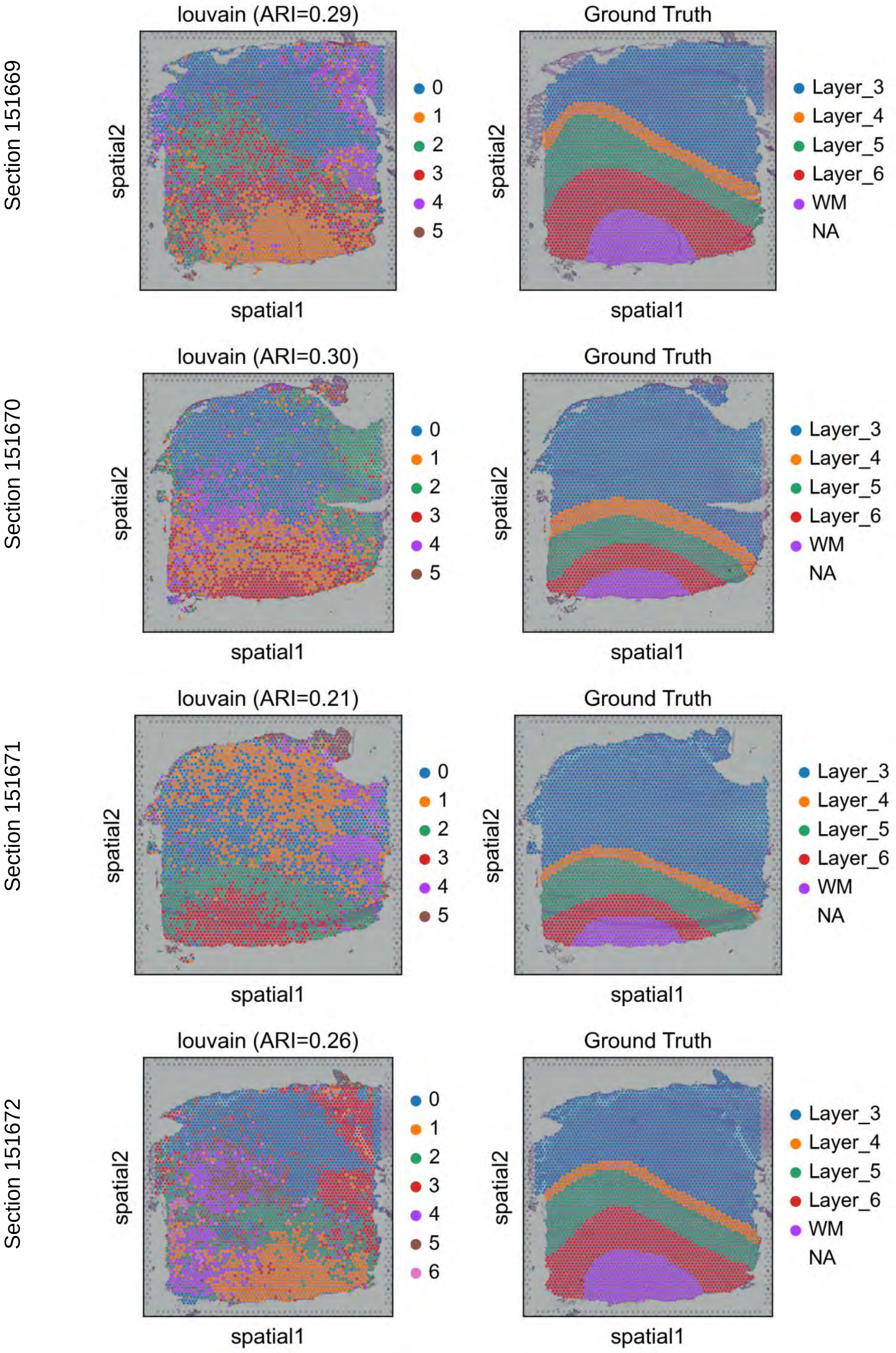


Supplementary Fig. S2. The result of Scanpy on DLPFC dataset sections 151569-151672.

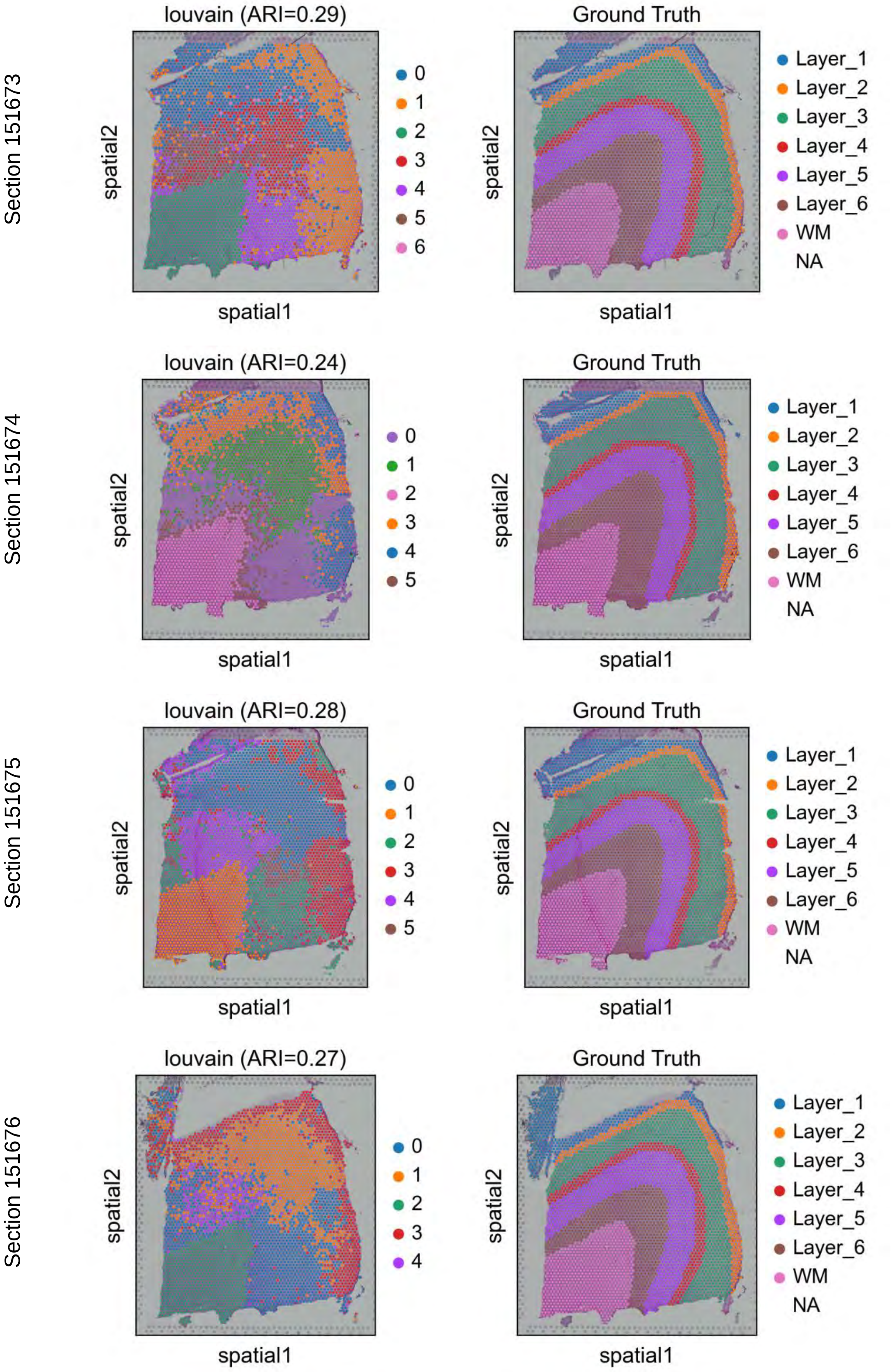


Supplementary Fig. S3. The result of Scanpy on DLPFC dataset sections 151573-151676.

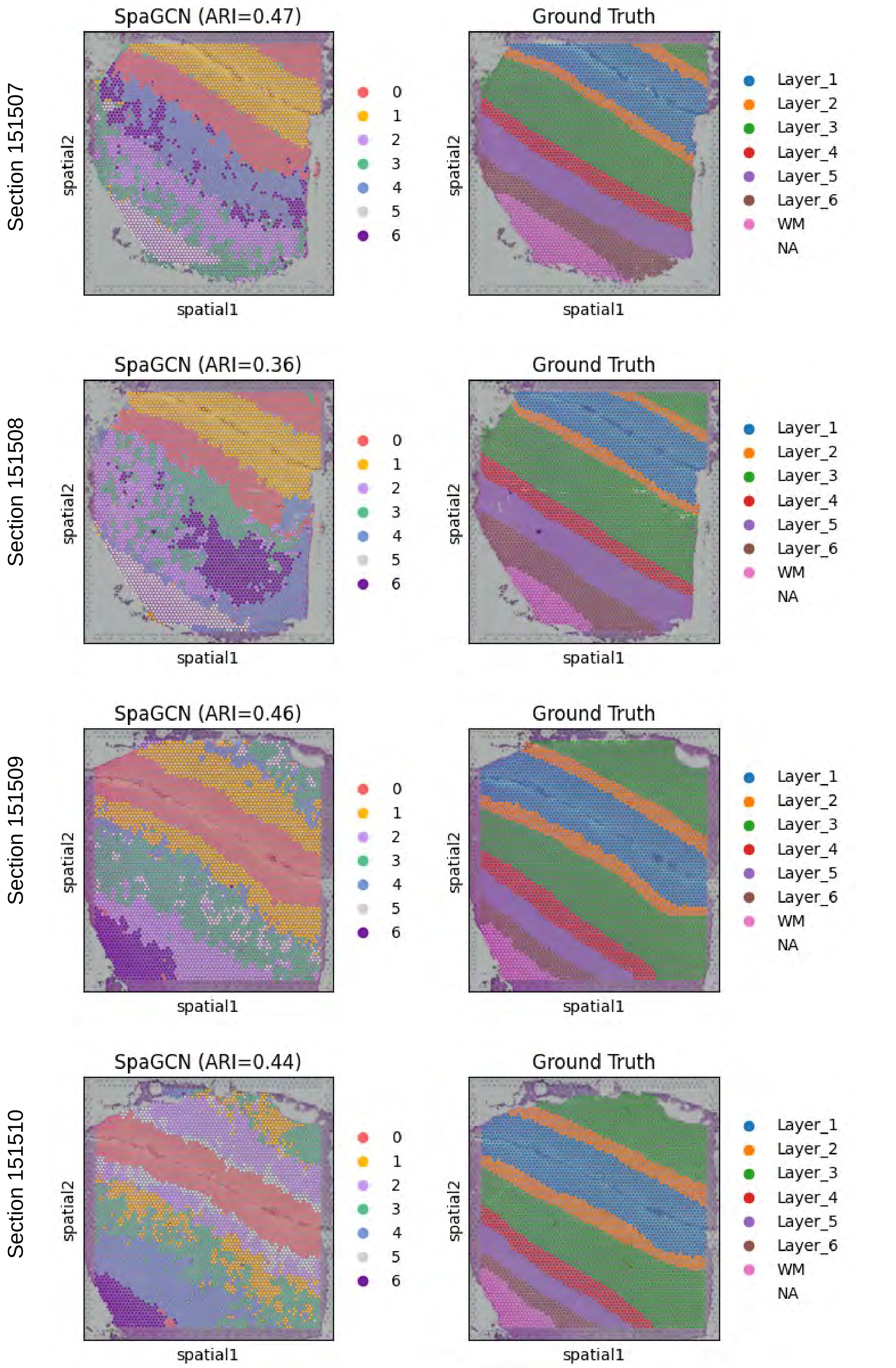


Supplementary Fig. S4. The result of SpaGCN on DLPFC dataset sections 151507-151510.

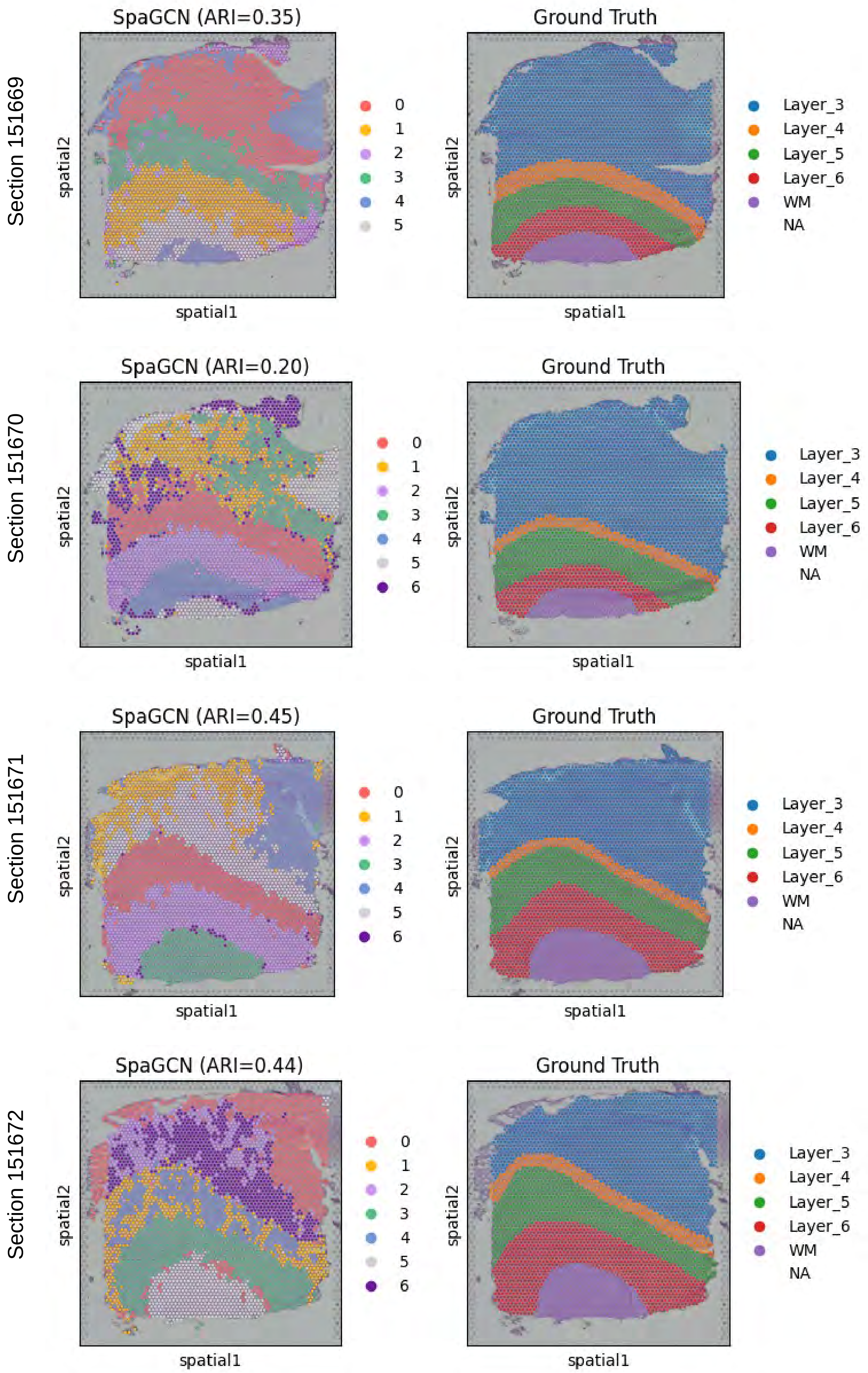


Supplementary Fig. S5. The result of SpaGCN on DLPFC dataset sections 151669-151672.

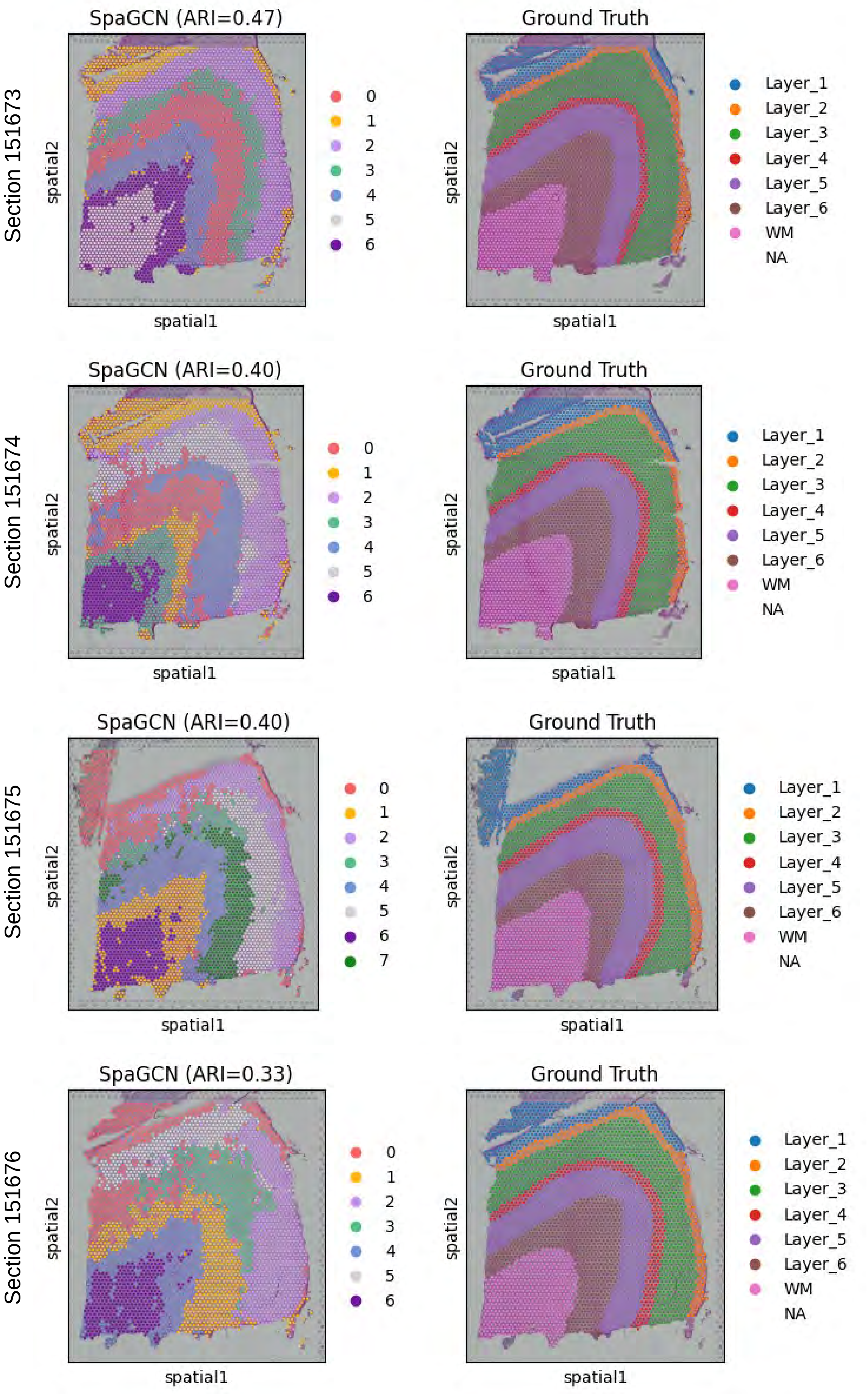


Supplementary Fig. S6. The result of SpaGCN on DLPFC dataset sections 151673-151676.

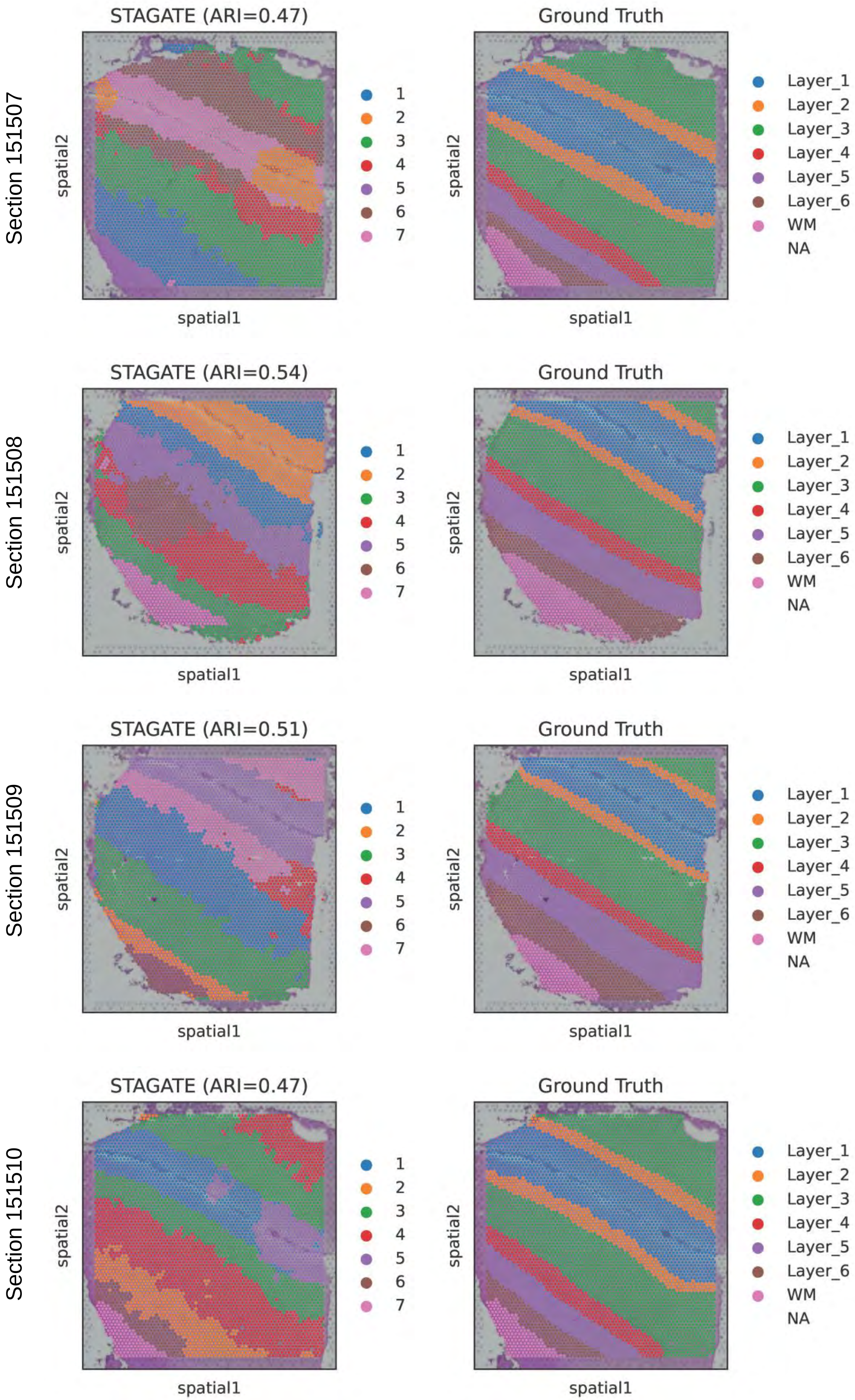


Supplementary Fig. S7. The result of STAGATE on DLPFC dataset sections 151507-151510.

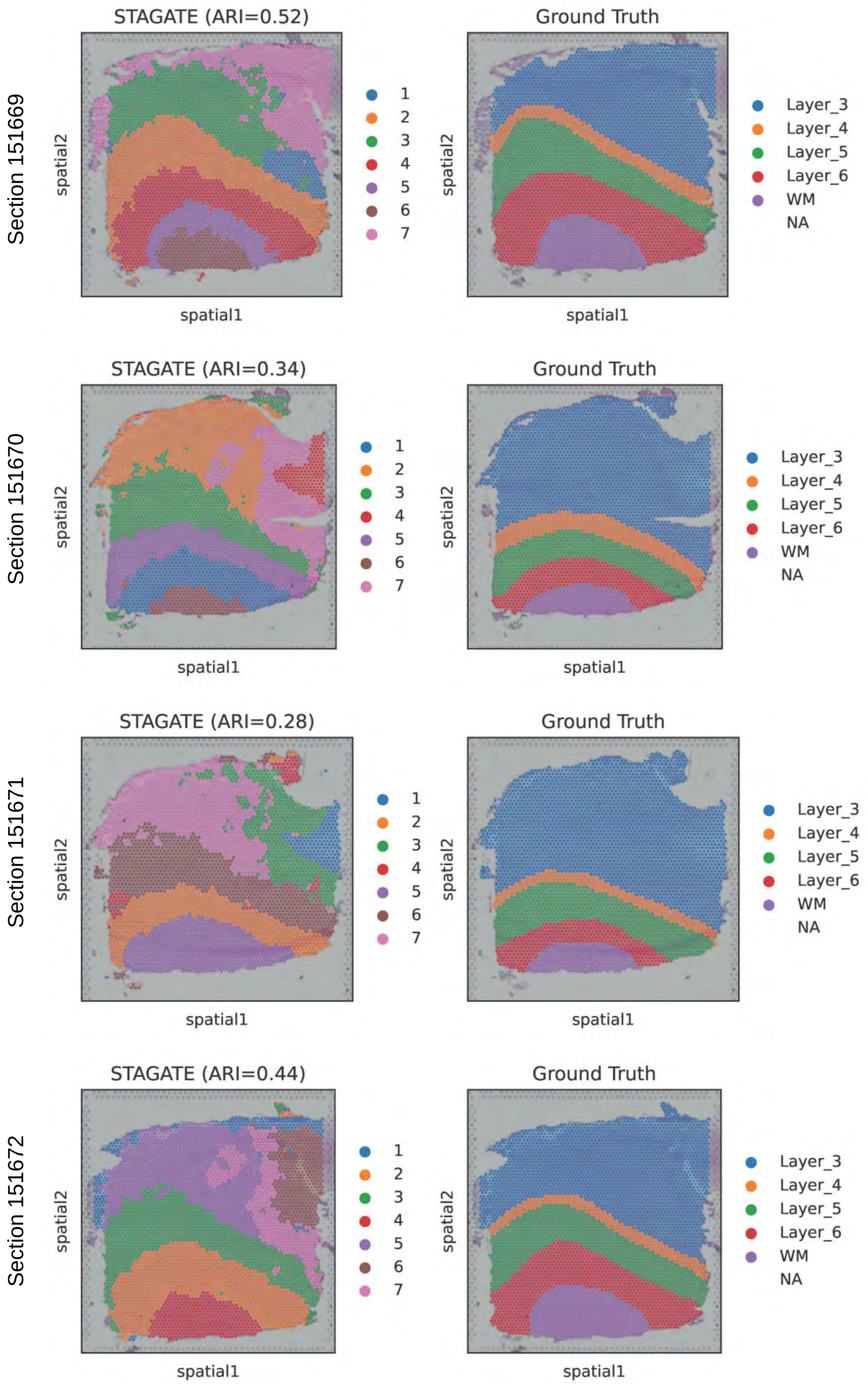


Supplementary Fig. S8. The result of STAGATE on DLPFC dataset sections 151669-151672.

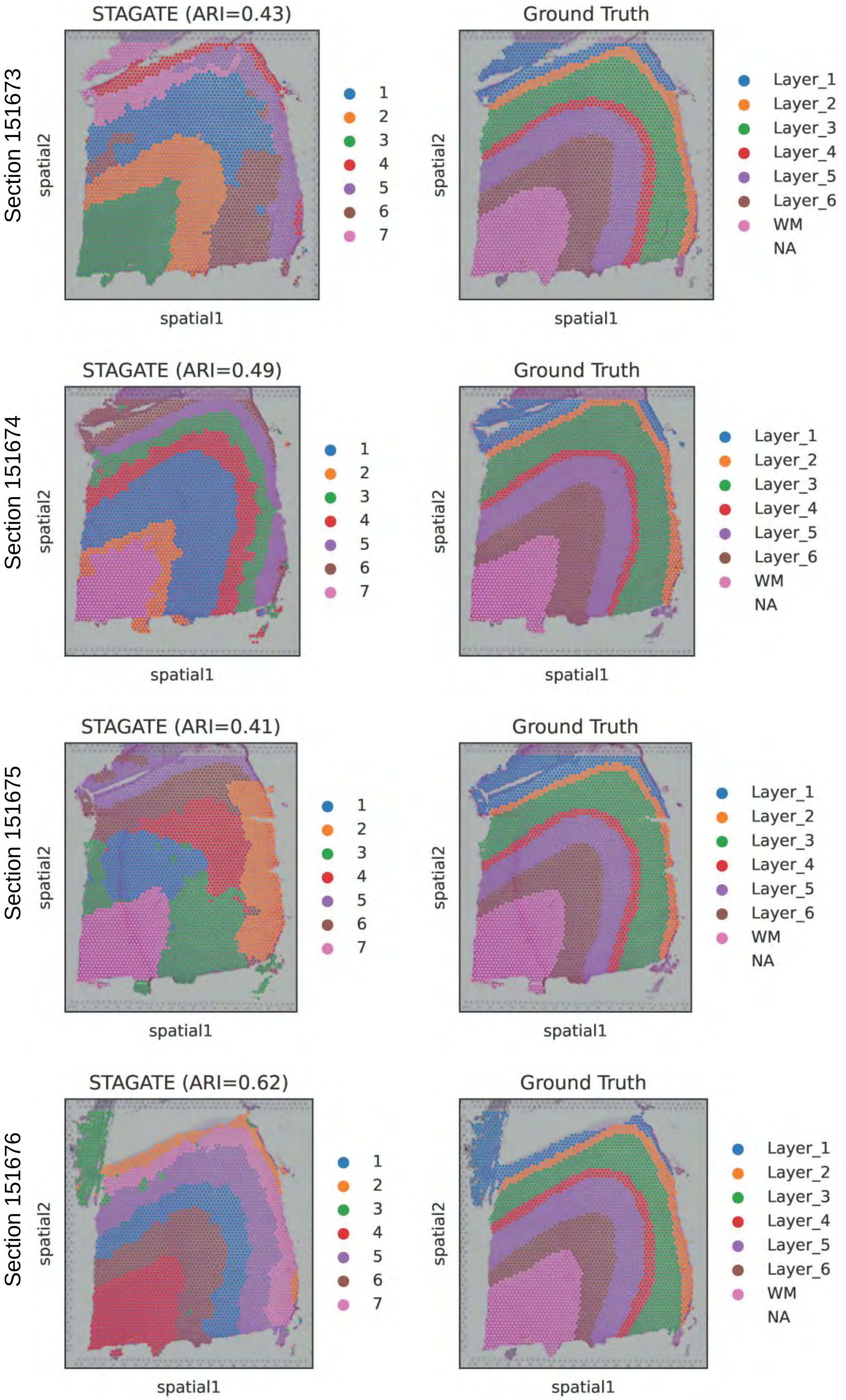


Supplementary Fig. S9. The result of STAGATE on DLPFC dataset sections 151673-151676.

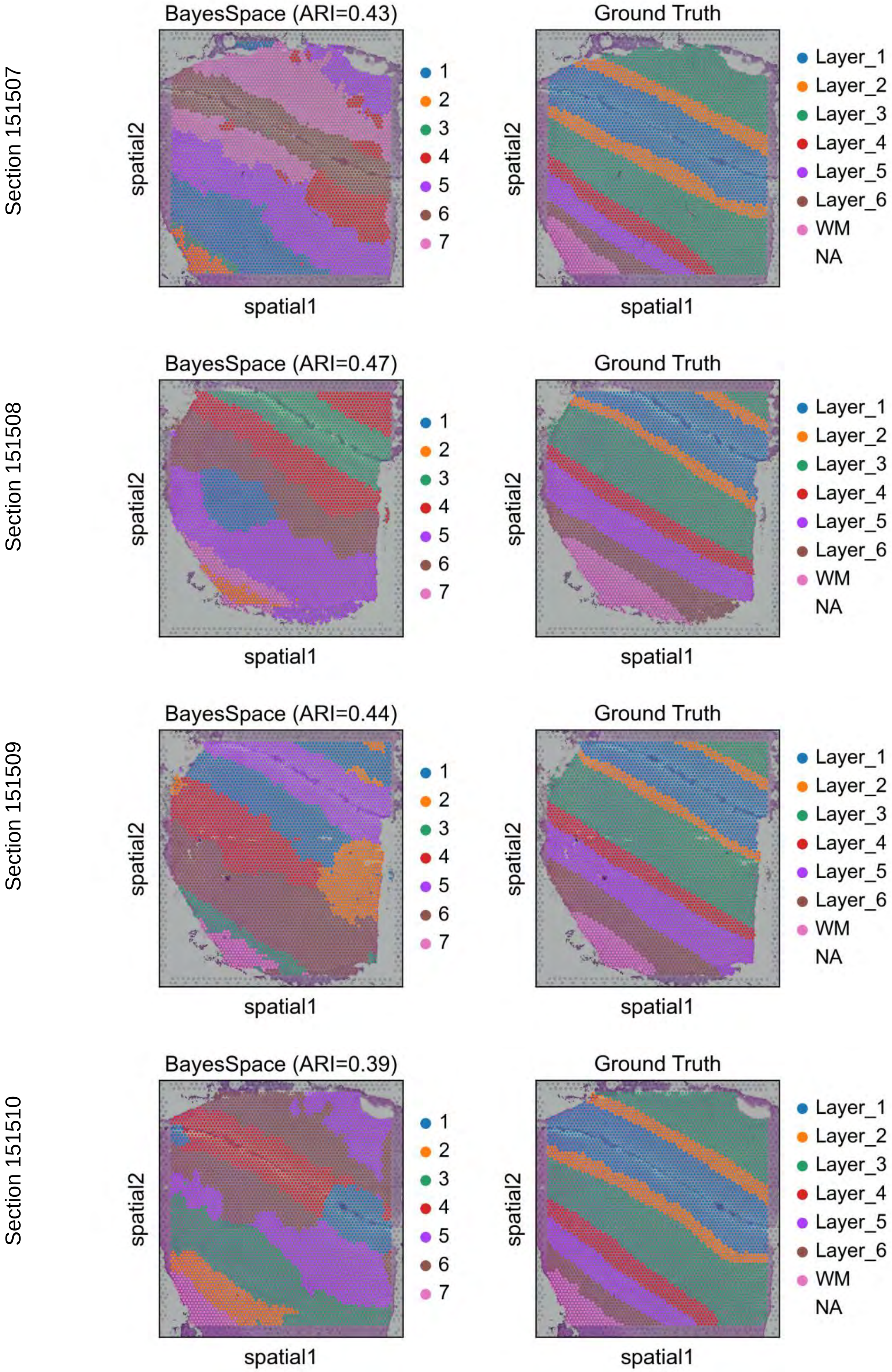


Supplementary Fig. S10. The result of BayesSpace on DLPFC dataset sections 151507-151510.

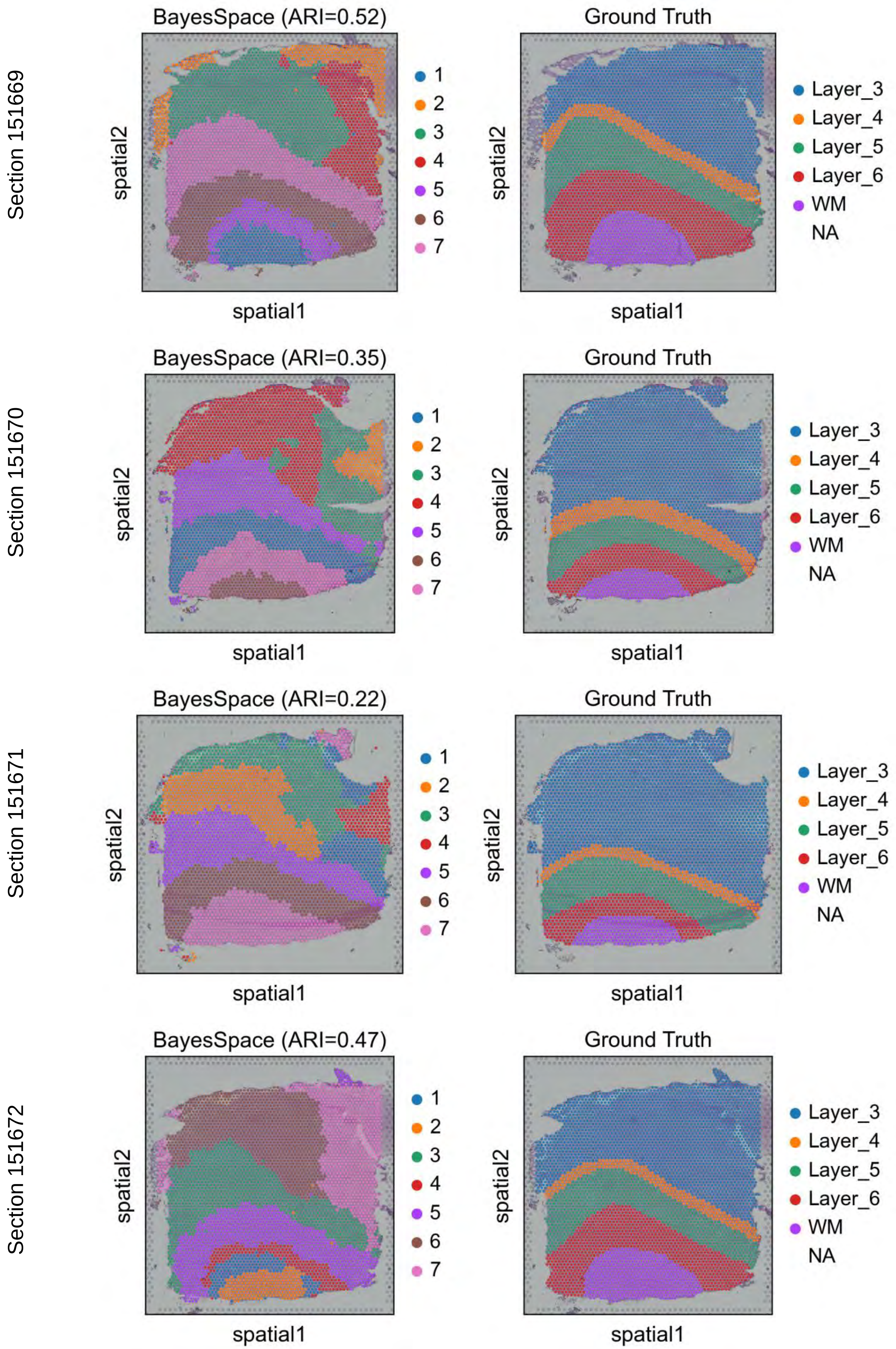


Supplementary Fig. S11. The result of BayesSpace on DLPFC dataset sections 151669-151672.

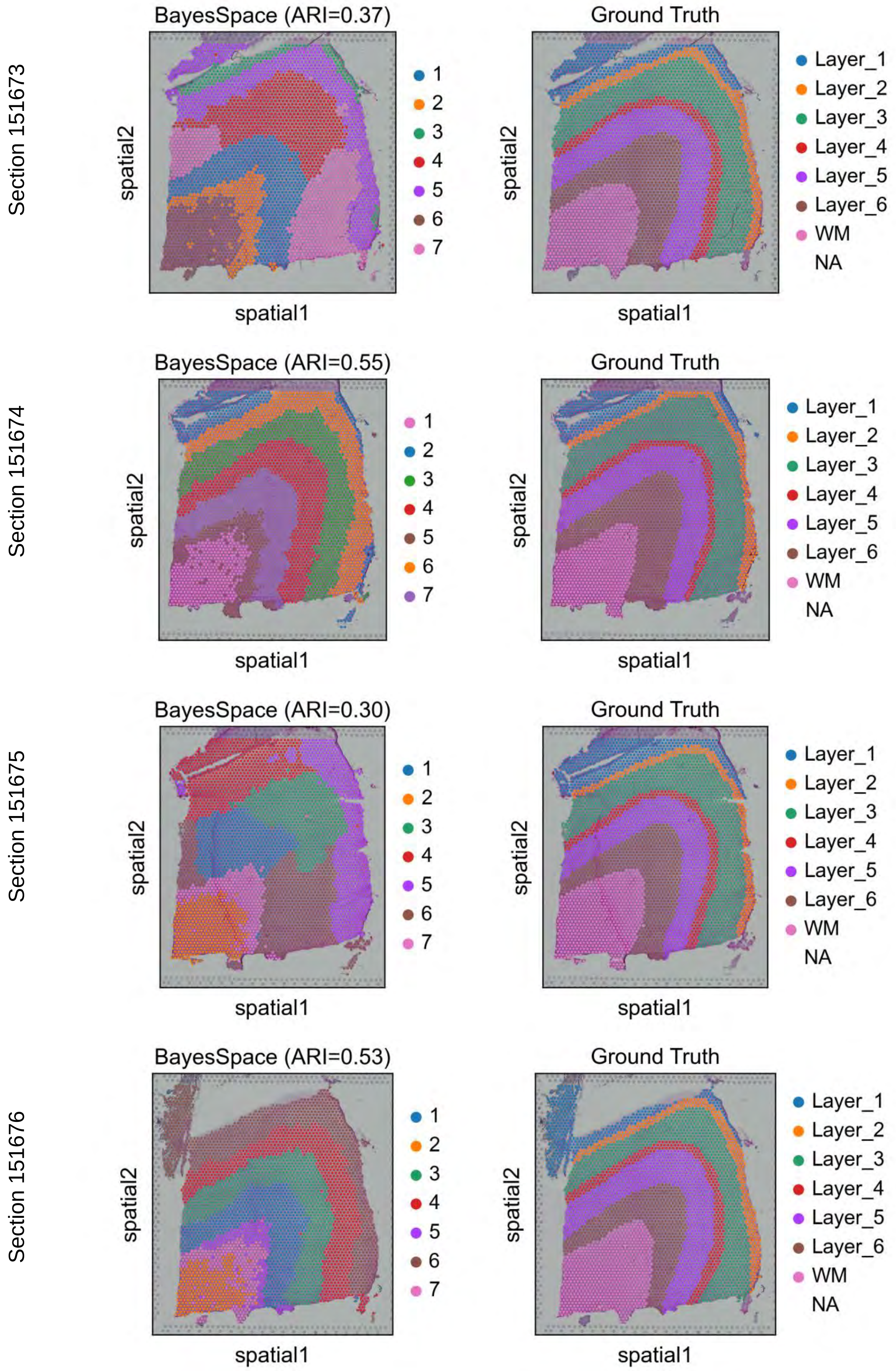


Supplementary Fig. S12. The result of BayesSpace on DLPFC dataset sections 151673-151676.

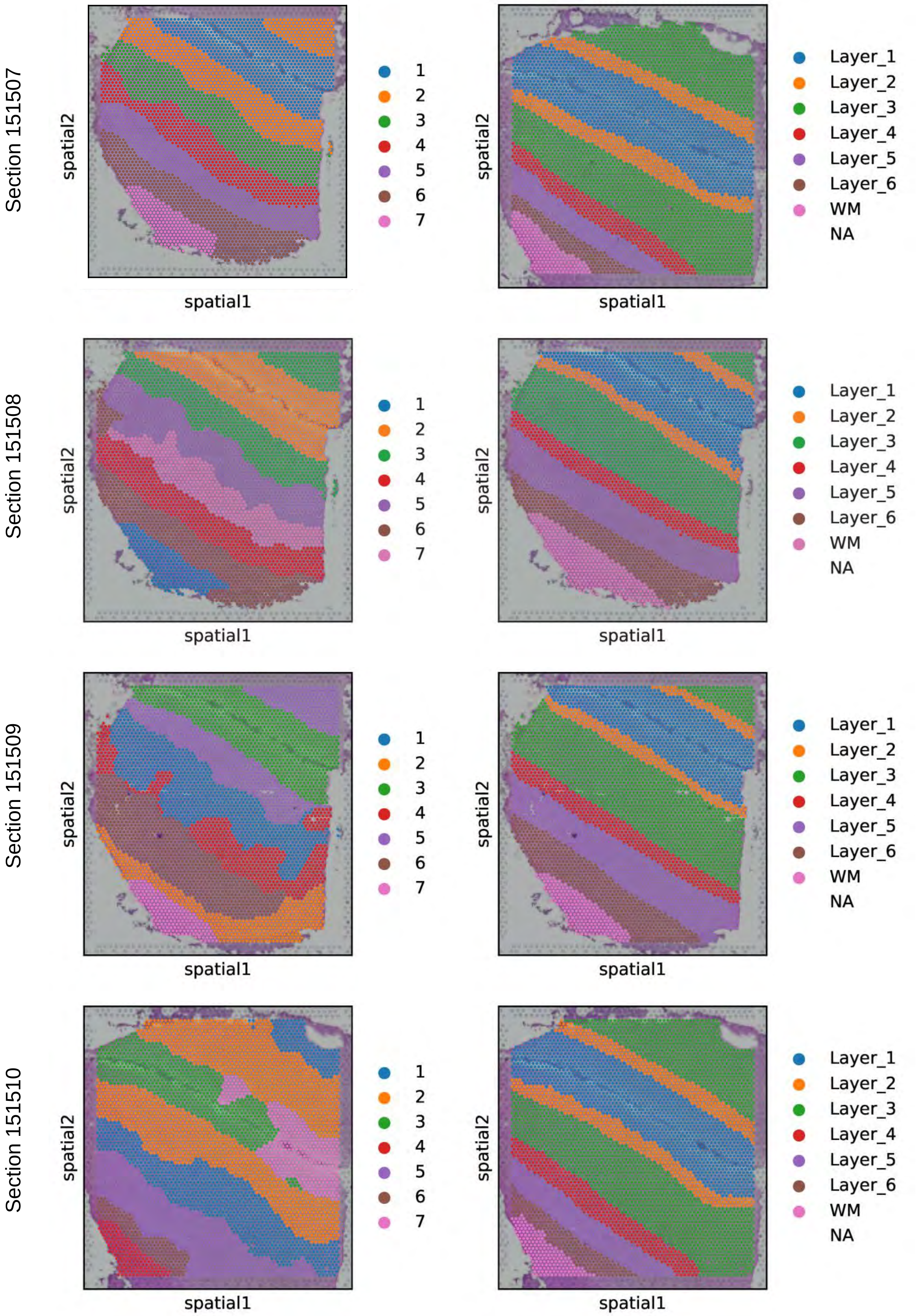


Supplementary Fig. S13. The result of HyperSTAR on DLPFC dataset sections 151507-151510.

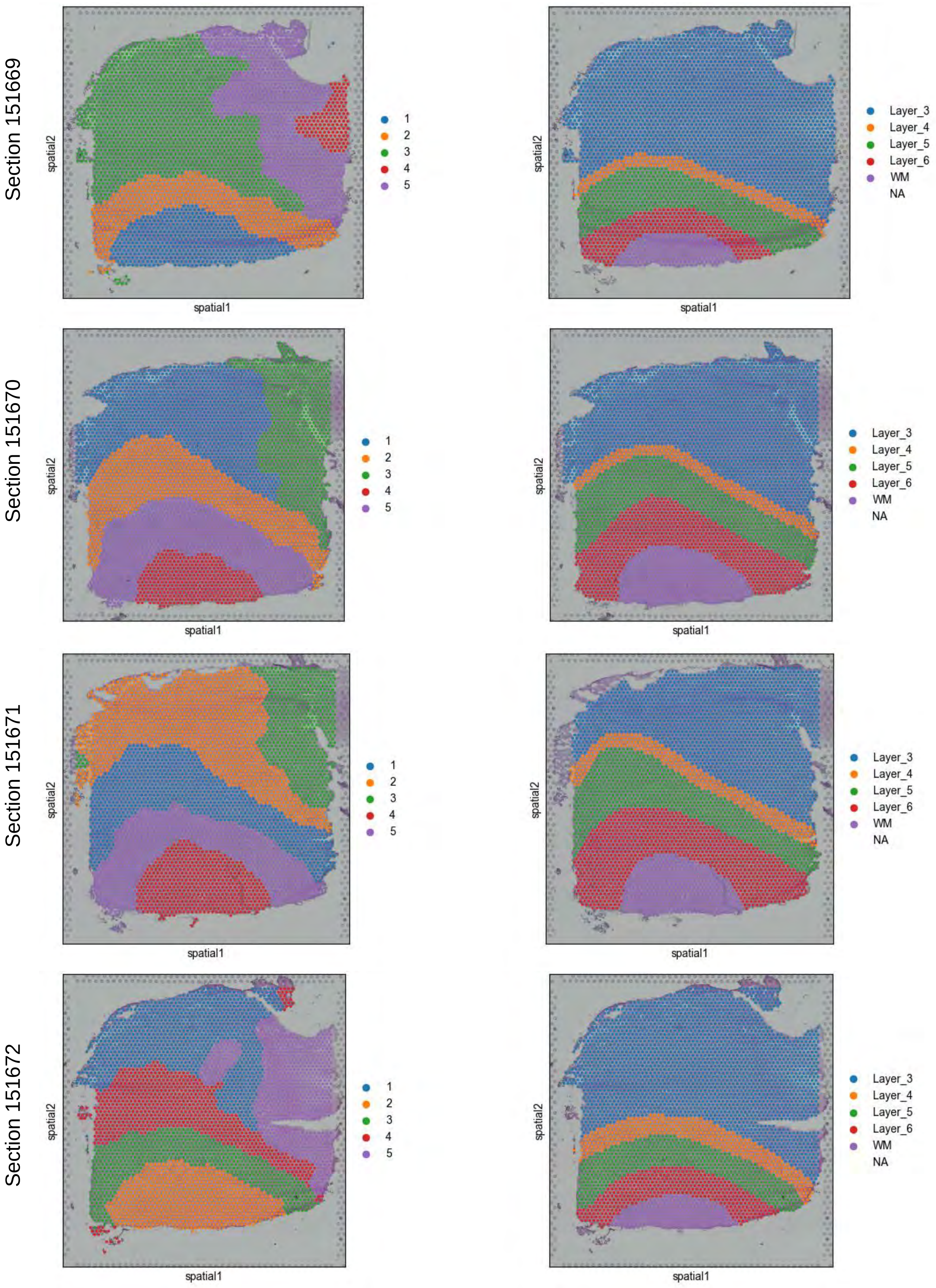


Supplementary Fig. S14. The result of HyperSTAR on DLPFC dataset sections 151669-151672.

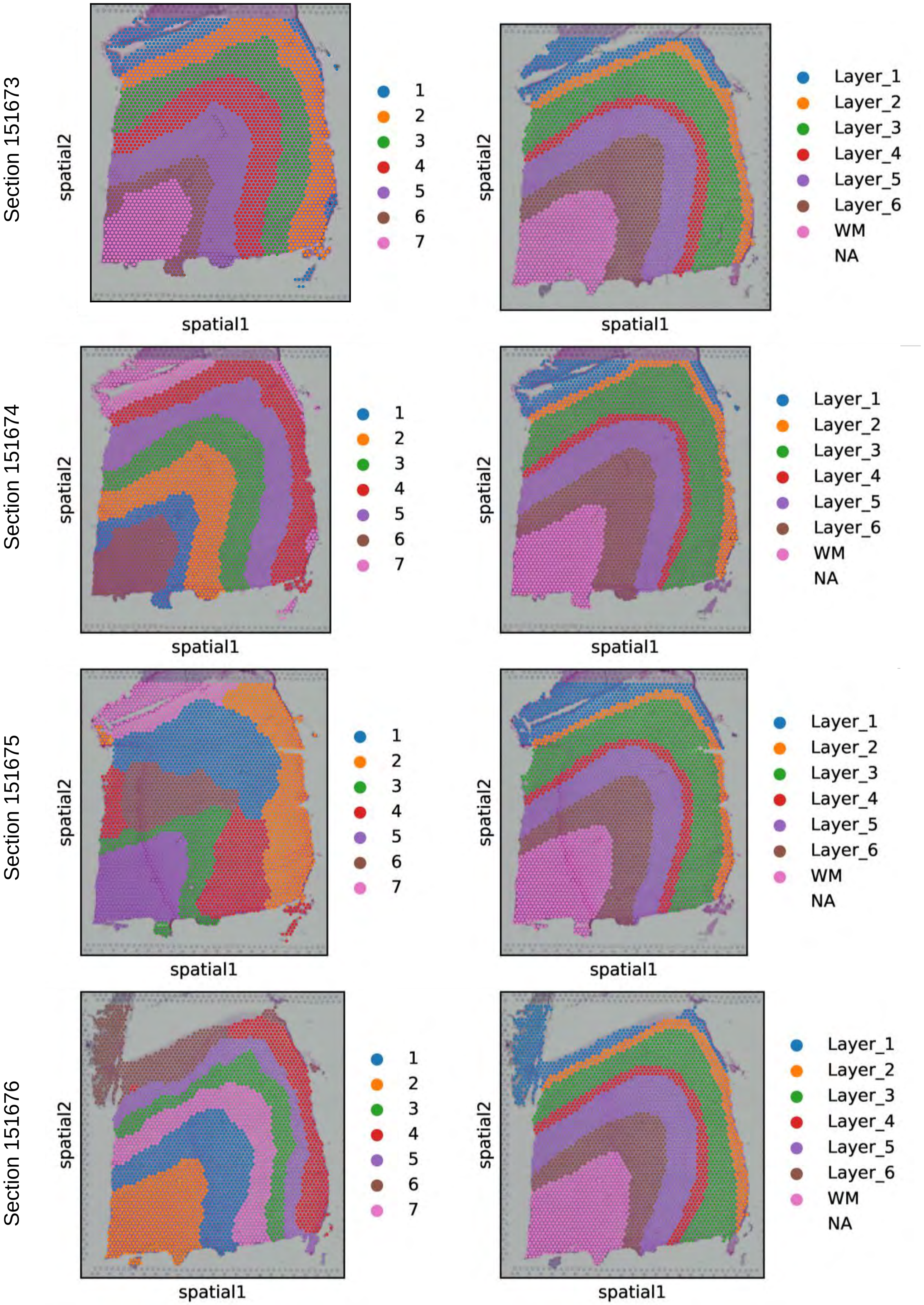


Supplementary Fig. S15. The result of HyperSTAR on DLPFC dataset sections 151673-151676.

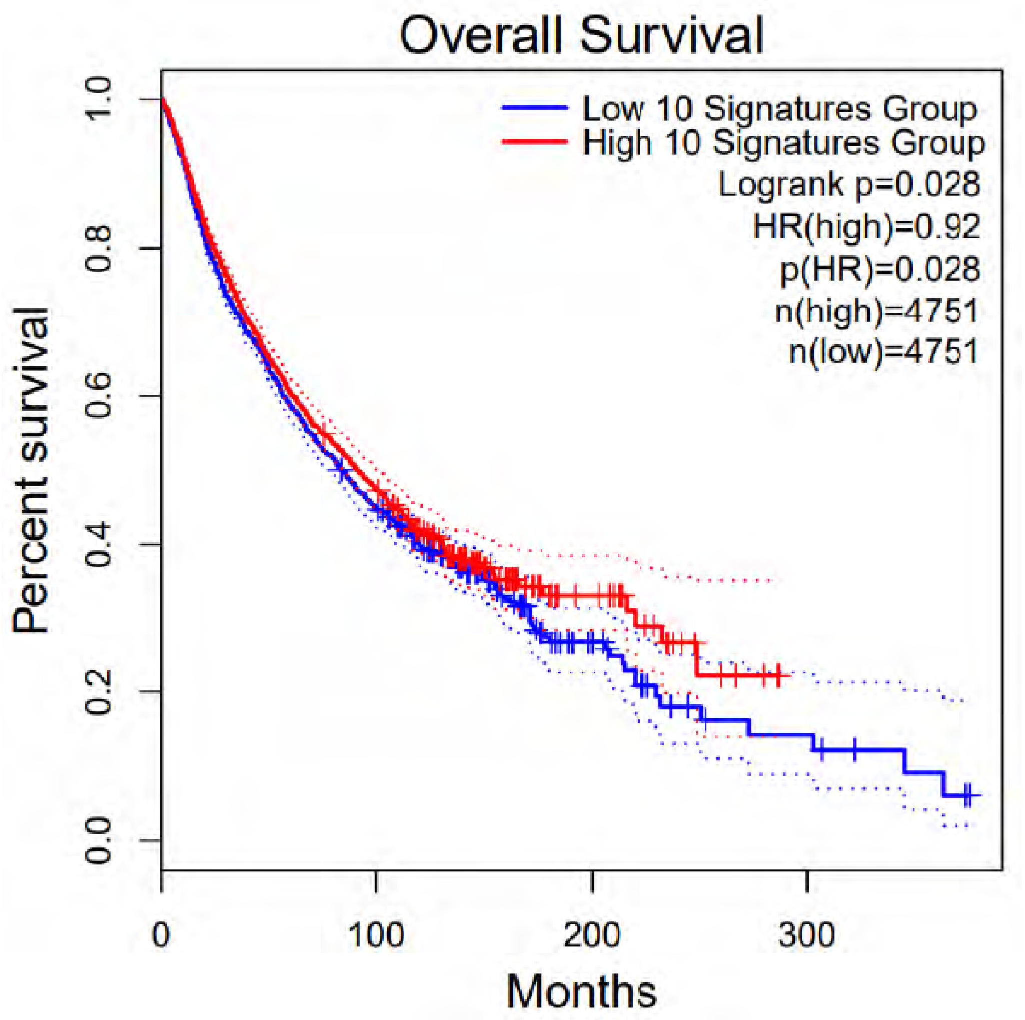


Supplementary Fig. S16. The survival analysis of IDC DEGs.

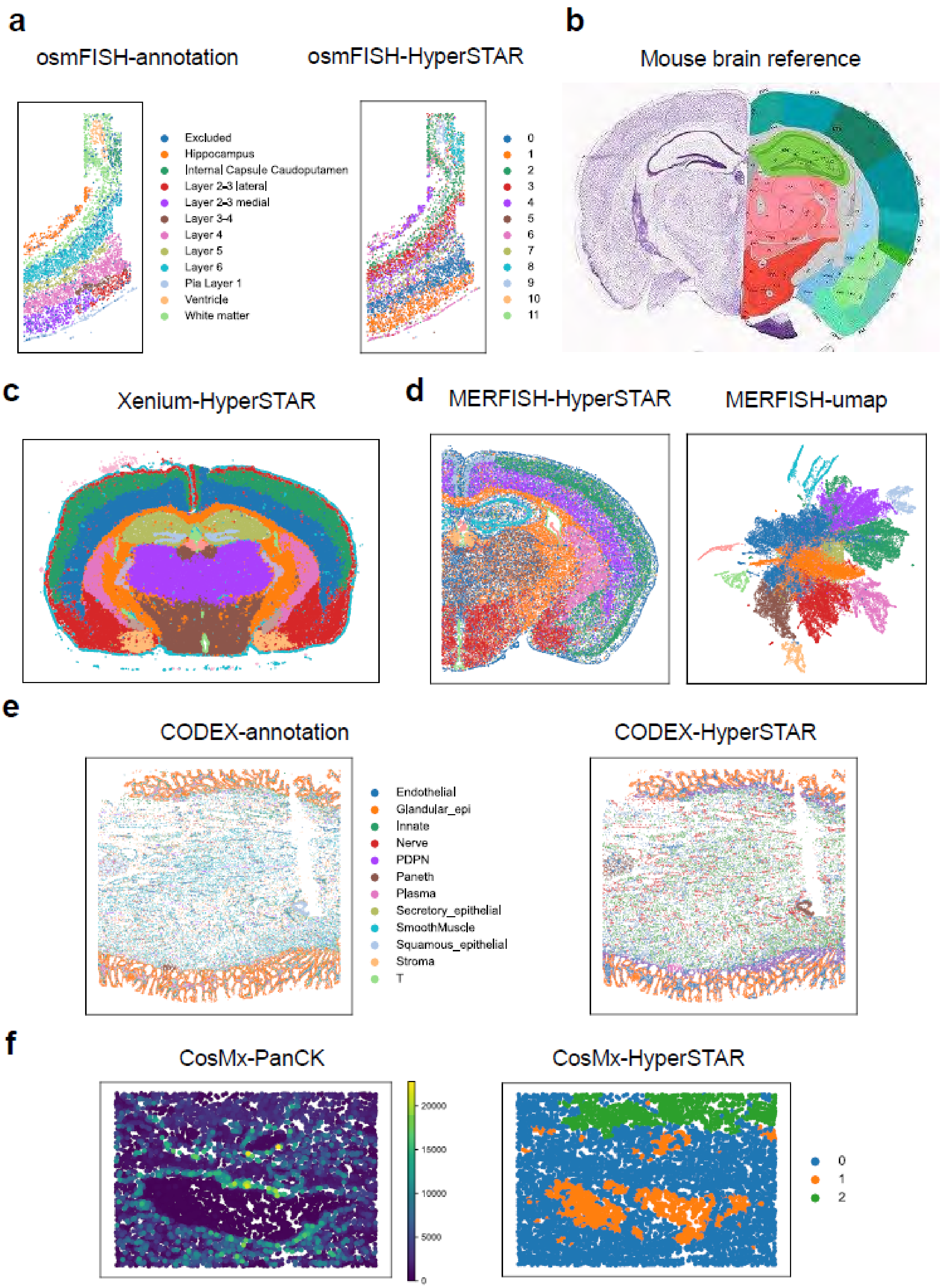
a
osmFISH-annotation
osmFISH-HyperSTAR
Excluded
Hippocampus
Internal Capsule Caudoputamen
Layer 2-3 lateral
Layer 2-3 medial
Layer 3-4
Layer 4
Layer 5
Layer 6
Pia Layer 1
Ventricle
White matter
0
1
2
3
4
5
6
7
8
9
10
11
b
Mouse brain reference
c
Xenium-HyperSTAR
d
MERFISH-HyperSTAR
MERFISH-umap
e
CODEX-annotation
CODEX-HyperSTAR
Endothelial
Glandular_epi
Innate
Nerve
PDPN
Paneth
Plasma
Secretory_epithelial
SmoothMuscle
Squamous_epithelial
Stroma
T
f
CosMx-PanCK
CosMx-HyperSTAR
20000
15000
10000
5000
0
0
1
2

Supplementary Fig. S17. HyerSTAR is applicable to various spatial omics and different platforms. **a** Manual annotation and HyperSTAR identification of somatosensory cortex from osmFISH. **b** The mouse brain reference from Allen Atlas. **c** The spatial result of HyperSTAR on a mouse brain section from Xenium. **d** The spatial result and UMAP plot by HyperSTAR on a mouse brain section from MERFISH. **e** Manual annotation and HyperSTAR identification of a Barrett's esophagus section from CODEX. **f** The PanCK distribution and HyperSTAR identification of a NSCLC section from CosMx.

**a**

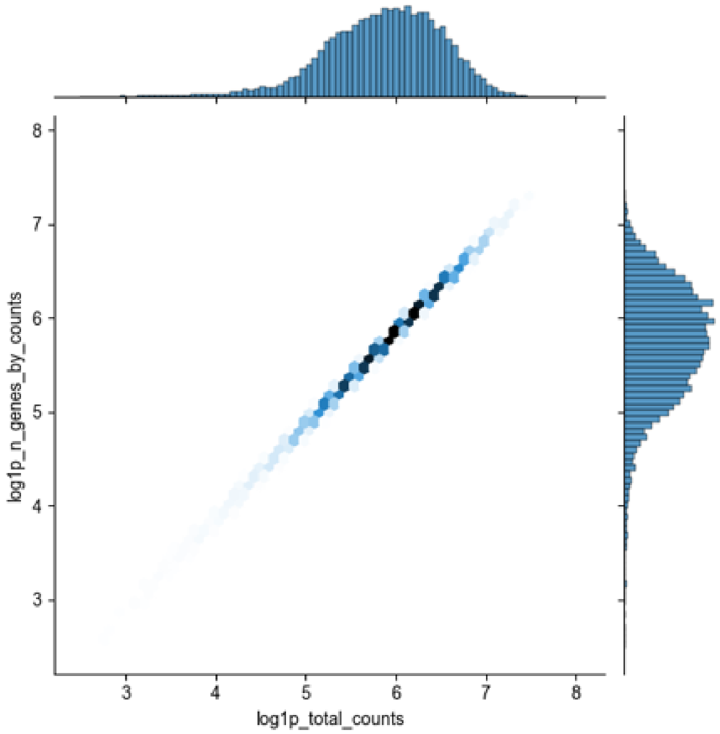


**b**

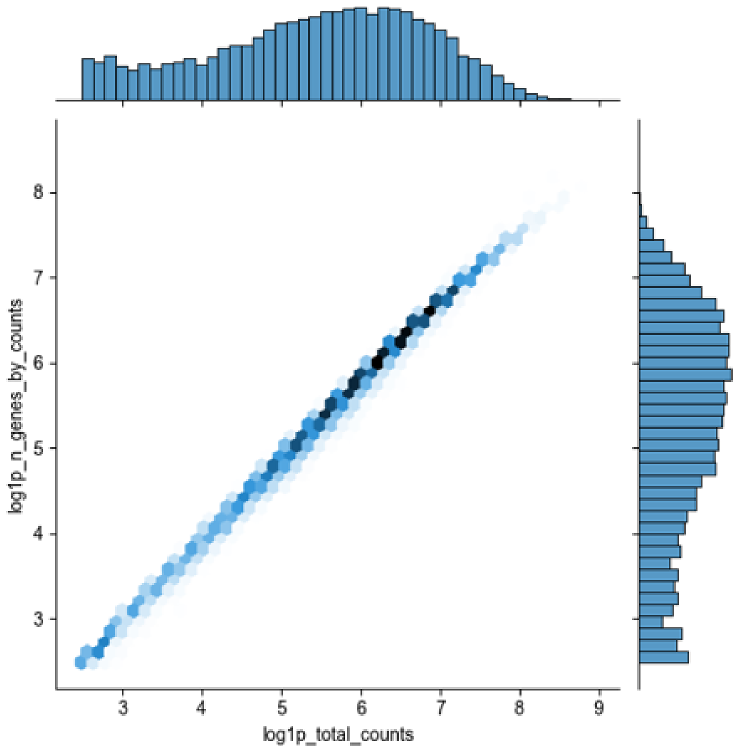


Supplementary Fig. S18. The data quality in mouse olfactory bulb data of Stereoseq and slideseqv2. **a** Data quality of Stereoseq data. **b** Data quality of slideseqV2 data.

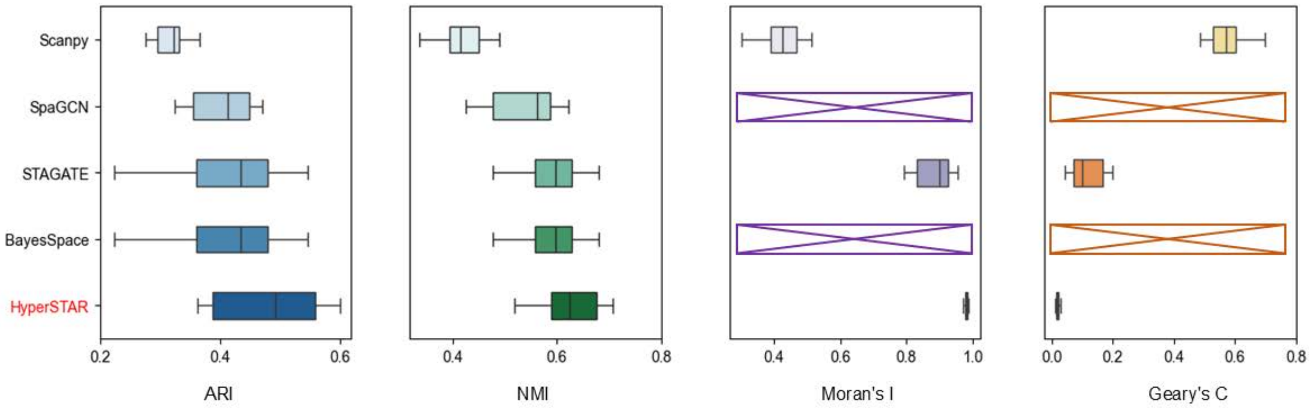


Supplementary Fig. S19. Boxplots comparing the performance of different methods across four metrics: ARI, NMI, Moran’s I, and Geary’s C. Each box represents the interquartile range (IQR) of the values for each method, with the median indicated by the line inside the box.

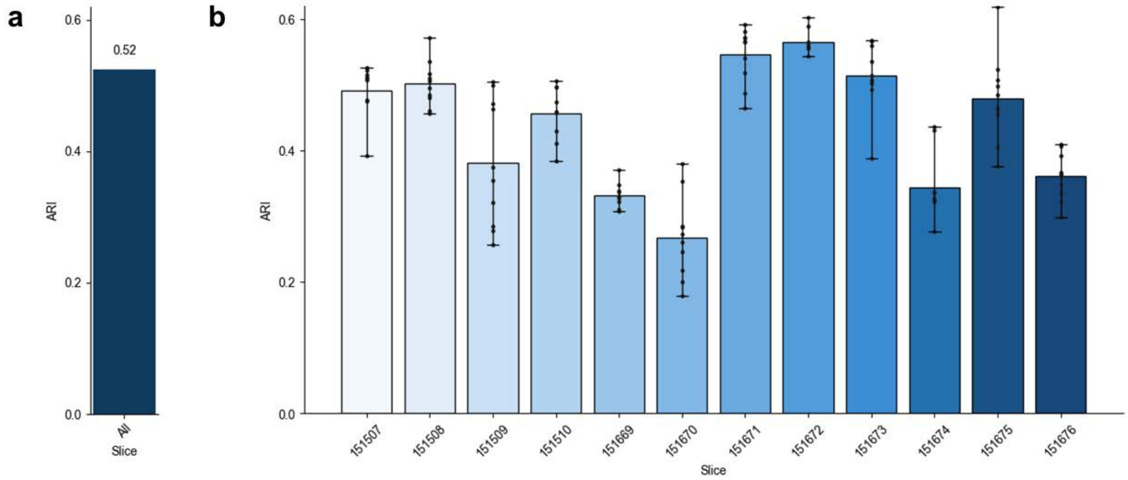


Supplementary Fig. S20. (a) The average Adjusted Rand Index (ARI) across 12 slices of the DLPFC dataset, with 10 independent runs for each slice. (b) ARI values obtained from 10 independent runs of HyperSTAR for each slice in the DLPFC dataset. The error bars represent the standard deviation of the ARI values across the 10 runs.

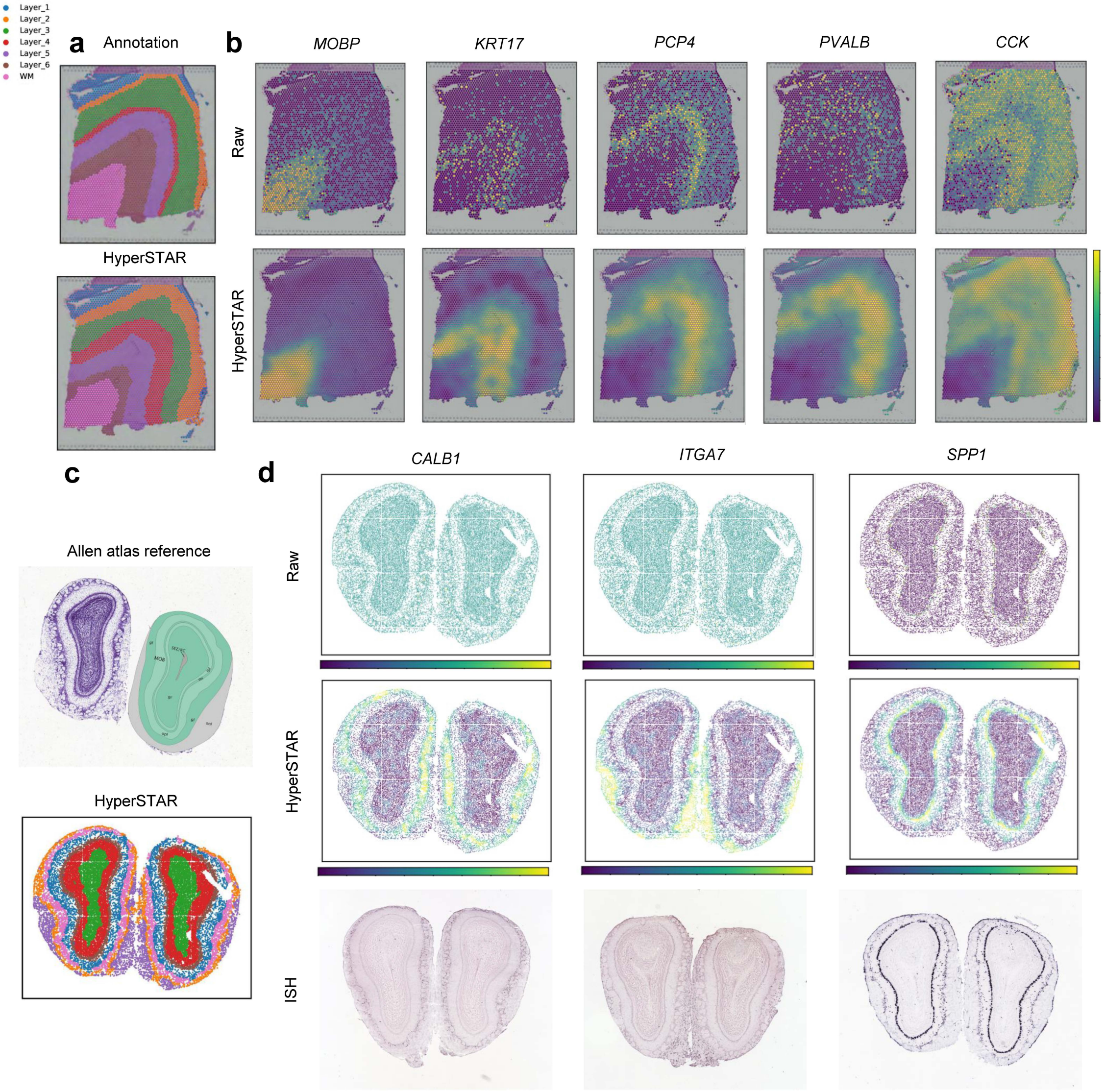


Supplementary Fig. S21. Analysis of Denoising Effects in Spatial Transcriptomics Data. (a) Annotation and HyperSTAR clustering results for DLPFC slice 151673. (b) Comparison of marker gene expression before and after denoising with HyperSTAR. (c) Allen reference annotations and HyperSTAR clustering for the mouse olfactory bulb. (d) Raw and HyperSTAR-denoised expression of marker genes alongside corresponding In Situ Hybridization (ISH) images from the Allen Brain Atlas.

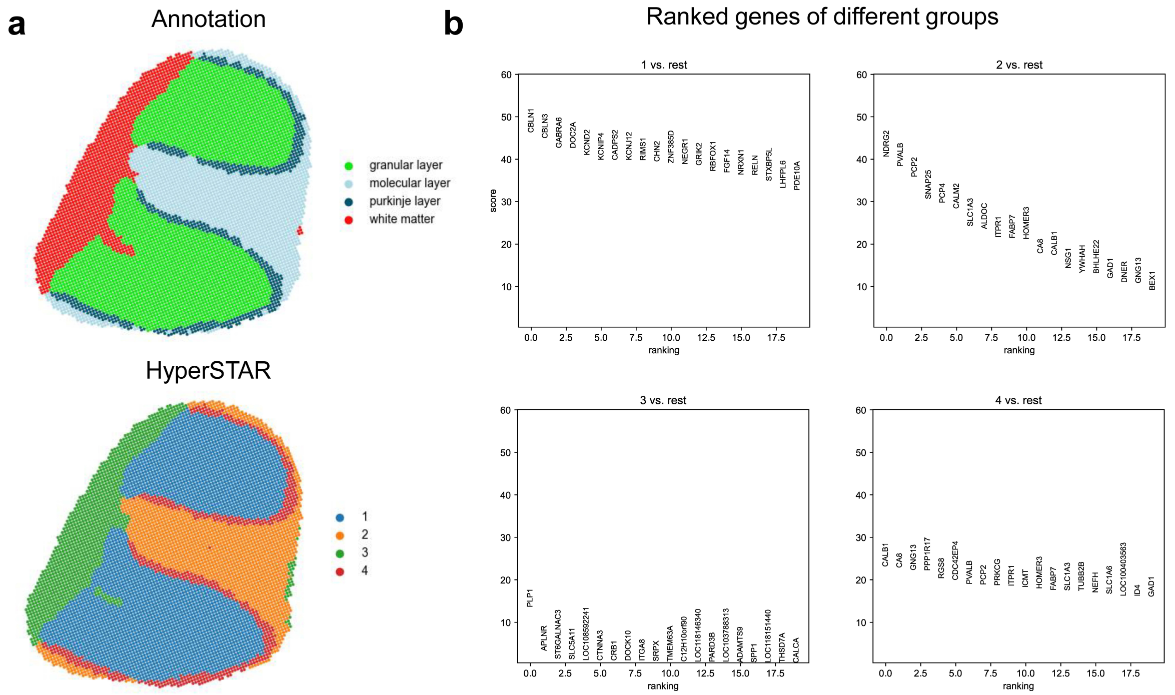


Supplementary Fig. S22. Identification of Spatially Variable Genes (SVGs) using HyperSTAR in the Marmoset Cerebellum. (a) Upper panel shows the expert annotations of the cerebellar sections, while the lower panel displays the clustering results obtained from HyperSTAR. (b) Visualization of the top Highly Variable Genes (HVGs) for each cluster identified by HyperSTAR, after applying sc.tl.rank_genes_groups().

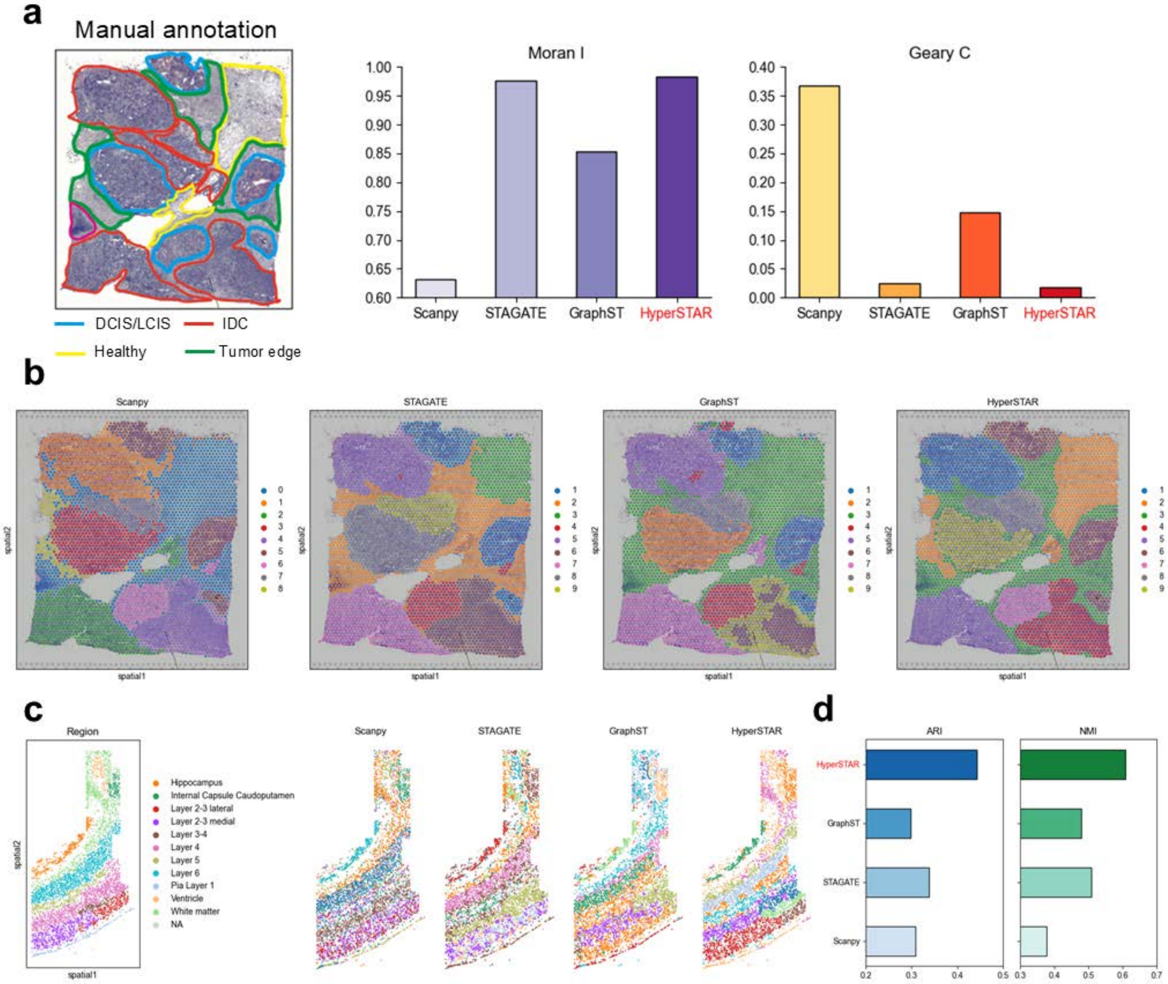


Supplementary Fig. S23. Comparative Performance of Clustering Methods on breast cancer and osmFISH dataset. (a) Manual annotations and metrics comparison for the breast cancer dataset, highlighting Moran's I and Geary's C. (b) Spatial visualization of clustering results from four methods (Scanpy, STAGATE, GraphST, HyperSTAR) on the breast cancer dataset. (c) Manual annotations alongside spatial visualizations of clustering outcomes for the osmFISH dataset using the same four methods. (d) Metrics comparison for the osmFISH dataset, including ARI and NMI.

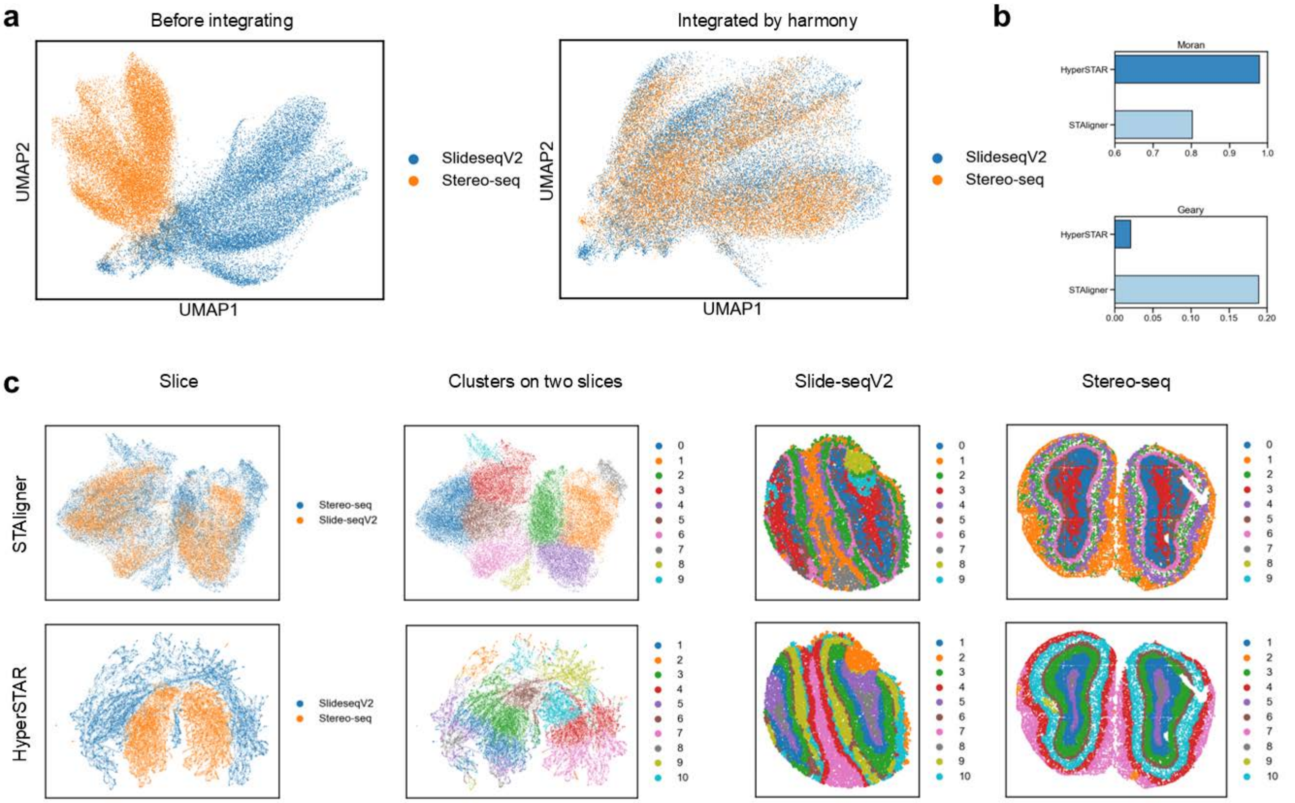


Supplementary Fig. S24. (a) UMAP plots depicting the data distribution before integration (left) and after Harmony integration (right). (b) Comparison of HyperSTAR and STAligner based on spatial autocorrelation metrics, Moran's I and Geary's C. (c) UMAP plots of the joint embedding (left) alongside spatial visualization of clustering results (right) for HyperSTAR (lower) and STAligner (upper).
